# Plant Kelch phosphatases are Ser/Thr phosphatases involved in cell cycle regulation

**DOI:** 10.64898/2026.01.06.697939

**Authors:** Felix Rico-Resendiz, Oded Pri-Tal, Pierre Raia, Andrea Moretti, Houming Chen, Jun Yu, Larissa Broger, Christelle Fuchs, Ludwig A. Hothorn, Sylvain Loubéry, Michael Hothorn

**Author notes:** current address: CSL Behring AG, 3014 Bern, Switzerland. These authors contributed equally to this work (Felix Rico-Resendiz, Oded Pri-Tal, Pierre Raia, Andrea Moretti). Corresponding author Michael Hothorn **Email:**.

## Abstract

Brassinosteroids (BRs) are plant steroid hormones sensed by the membrane receptor kinase BRI1. Activation of BRI1 leads to the dephosphorylation of BZR1/BES1 transcription factors. Overexpression of the Kelch phosphatase BRI1 SUPPRESSOR 1 (BSU1) rescued the growth defects of *bri1* mutants. Subsequent studies identified BSU1 as a protein tyrosine phosphatase, which promotes BR signaling by dephosphorylating a phosphotyrosine in the glycogen synthase kinase 3 BIN2. Crystal structures of the BSU1 phosphatase domain now reveal a high degree of structural similarity to protein phosphatase 1 (PP1), a eukaryotic serine/threonine phosphatase. Consistently, BSU1 efficiently dephosphorylated phosphothreonine– and phosphoserine-containing substrate peptides, but showed no detectable activity toward BIN2 and other phosphotyrosine substrates. A catalytically inactive BSU1 phosphatase domain suppresses the growth phenotypes of the Arabidopsis *bri1-5* mutant and binds the BSU1 homologs BSL1-3. *bsu1* and *bsu1 bsl1 bsl2/3* loss-of-function mutants display wild-type-like BR responses, but exhibit stomatal patterning and fertility defects. Importantly, the PP1-like C-terminal tail of BSU1 is phosphorylated at Thr785 by a cyclin-dependent kinase complex. The phosphorylated tail binds to the BSU1 substrate-binding grooves, blocking access to the active site. Mutation of Thr785 to alanine activates BSU1, suggesting that Kelch phosphatases and PP1 share a common regulatory mechanism. Deletion of the *Marchantia polymorpha* Kelch phosphatase MpBSLM results in an undifferentiated cell mass phenotype, associated with the over-activation of a cell cycle reporter. Taken together, our experiments suggest that plant Kelch phosphatases act as PP1-like cell cycle regulators, rather than as tyrosine phosphatases in brassinosteroid signaling. (243 words)

**Significance Statement:** Brassinosteroid hormones initiate a well-characterized signaling pathways in plants. While the early signaling events at the plasma membrane have been thoroughly examined, the precise sequence of molecular events that comprise the cytoplasmic signaling cascade is less well-understood. Here, the molecular function of the Kelch protein phosphatase BSU1 is re-evaluated. Previous reports have indicated that BSU1 dephosphorylates the GSK3 kinase BIN2 on a central tyrosine residue. However, structural, quantitative enzyme kinetic, and genetic analyses now suggest that BSU1 is a PP1-like serine/threonine phosphatase involved in cell cycle control rather than a tyrosine phosphatase involved in BIN2 de-phosphorylation. The regulation of BSU1 by a cyclin-dependent kinase complex and the role of Kelch phosphatases in cell cycle progression in *Marchantia polymorpha* are experimentally characterized. (121 words)

## Introduction

Brassinosteroids (BRs) are a class of steroid hormones (1, 2) that regulate plant growth and development by binding to the membrane receptor kinase BRASSINOSTEROID INSENSITIVE 1 (BRI1) (3–6). BR binding to the BRI1 leucine-rich repeat extracellular domain (7, 8) promotes the dissociation of SOMATIC EMBRYOGENESIS RECEPTOR KINASES (SERKs) from receptor pseudokinases (9, 10), and the formation of a BRI1 – BR – SERK signaling complex (11–15, 6). The ligand-induced interaction of receptor and co-receptor at the cell surface brings their cytoplasmic kinase domains in proximity, triggering transphosphorylation and activation of the BRI1 kinase domain (16–19). The exact sequence of the early cytoplasmic signaling events has not been defined. The inhibitor protein BKI1 dissociates from the BRI1 kinase domain following receptor activation and relocates from the plasma membrane to the cytosol (20–22), where it associates with 14-3-3 proteins (23, 24). The BRI1 kinase domain can associate with different substrates, including the plasma membrane proton pump (25) and several cytoplasmic signaling kinases. BRI1-SIGNALING KINASES (BSKs) (26) form a family of protein kinases / pseudokinases with a conserved kinase core and a C-terminal tetratricopeptide repeat (TPR) domain (27). BSK family members are phosphorylated by BRI1 on a conserved serine residue (Ser230 in BSK1) in the activation segment of the kinase domain upon BR treatment in vivo (26, 28), and constitutively in vitro (26, 29, 24). Consistent with a function in BR signal transduction, *bsk* loss-of-function alleles have reduced BR signaling responses and associated growth phenotypes (26, 29–31). A second set of plasma-membrane associated, cytosolic BRI1 substrates are the protein kinases CONSTITUTIVE DIFFERENTIAL GROWTH 1 (CDG1) and CDL1 (32, 33). Both BSKs and CDG1/CDL1 have been reported to interact with the protein phosphatase BRI SUPPRESSOR 1 (BSU1) (33, 34). BSU1 was initially identified as a dominant suppressor of the *bri1-5* BR-receptor allele in an activation tagging (35) forward genetic screen (36). BSU1 contains an N-terminal Kelch repeat and a C-terminal phosphatase module, a domain architecture only found in plants and some unicellular eukaryotes (37, 38) (*SI Appendix*, Fig. S1). In Arabidopsis, there are three BSU1 homologs (BSL1-3) and over-expression of *BSU1*, *BSL1* and *BSL2* partially rescued the growth and BR signaling defects of both weak and strong BR-receptor alleles (33, 34, 36, 39, 40). *bsu1* T-DNA insertion mutants had no obvious phenotype (36, 37), while *bsl2 bsl3* mutants were small, sterile, and had short roots and twisted leaves (37). No homozygous *bsl1 bsl2 bsl3* mutants could be obtained (37). However, *bsu1 bsl1 BSL2*/*BSL3*-amiRNA plants had strongly reduced BR responses and an associated dwarf growth phenotype (34). At the biochemical level, both BSK1 and CDG1/CDL1 interacted with BSU1 in co-immunoprecipitation assays, in yeast two-hybrid experiments, and in pull downs (33, 34, 36, 30). CDG1/CDL1 phosphorylated BSU1 Ser 764 in vitro. BSU1 in turn dephosphorylated the glycogen synthase kinase 3 (GSK3) BRASSINOSTEROID-INSENSITIVE 2 (BIN2) (41) at Tyr200, a GSK3 autophosphorylation site important for enzyme function (42, 43), in vitro (33, 34). Together, these experiments defined BSU1 as a tyrosine phosphatase that inactivates BIN2, thereby decreasing BIN2-mediated phosphorylation of the BR transcription factors BZR1 and BES1, ultimately modulating BR-responsive gene expression (44–50). Outside BR signaling, plant Kelch phosphatases have been characterized as components of mitogen-activated protein kinase signaling pathways regulating stomatal development (51–53) or immune signaling (54–56) in Arabidopsis, and in cell cycle progression in *Chlamydomonas reinhardtii* (57, 58).

We aim to dissect the molecular mechanism of brassinosteroid signaling from BR sensing at the plasma membrane to the activation of nuclear transcription factors (59). Here we present the structure and enzyme mechanism of BSU1 and (re-)investigate its function in BR and cell signaling in Arabidopsis and in *Marchantia polymorpha*.

## Results

### BSU1 is a PP1-like Ser/Thr phosphatase

We expressed BSU1 in baculovirus-infected insect cells (*SI Appendix*, Fig. S2) and obtained crystal structures of its N-terminal Kelch (residues 11-351) and C-terminal phosphatase (residues 450-793) domains at 2.25 and 2.1 Å resolution, respectively (Fig. 1 *A*-*C*) (*SI Appendix*, Table S1). The Kelch domain is composed of a 6-bladed β-propeller (Fig. 1*B*), which shares structural similarity with many Kelch domain-containing proteins mediating protein-protein interactions, including human E3 ubiquitin ligases (60, 61) (*SI Appendix*, Fig. S3 *A* and *B*). Ser251, which is phosphorylated by the immune signaling kinase BIK1 (54), maps to a loop connecting blade IV and V (Fig. 1*B*). The C-terminal phosphatase domain has a canonical protein phosphatase fold (Fig. 1*C*) (*SI Appendix*, Fig. S3*C*). An extended C-terminal tail folds back into the active site of the enzyme and is well-defined in our structure (Fig. 1*C*). Ser764, which is phosphorylated by CDG1/CDL1, maps to the C-terminal tail of BSU1 (Fig. 1*C*) (*SI Appendix*, Fig. S1).

**Fig. 1.**
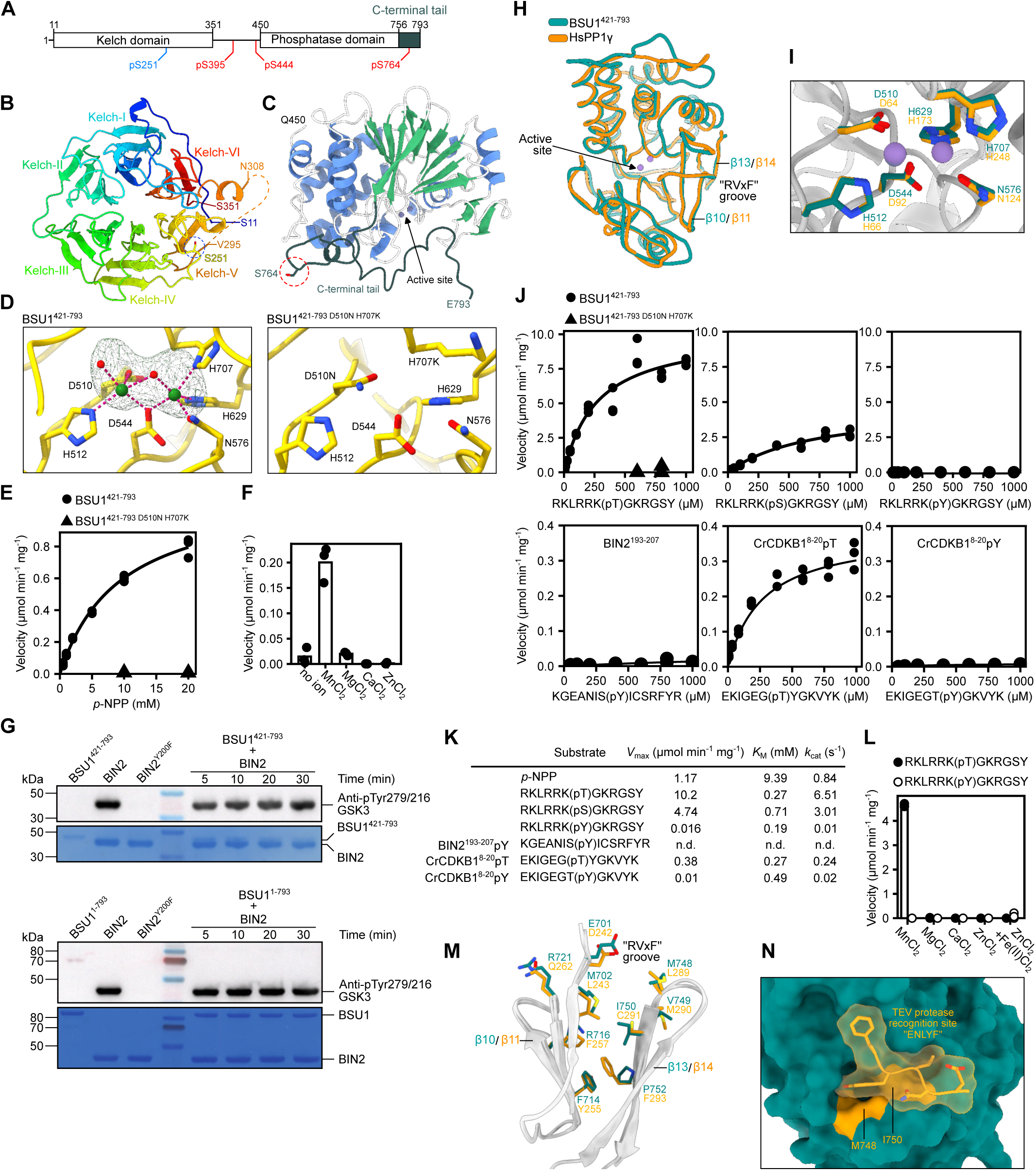
BSU1 is a PP1-like Ser/Thr phosphatase. (*A*) Schematic representation of *A. thaliana* BSU1, depicting the Kelch (residues 11–351) and phosphatase (residues 450–793) domains (https://www.uniprot.org/ Uniprot ID Q9LR78). Structured regions are shown as blocks, putative linker regions (residues 1–10 and 352–449) as solid line, CDG1 phosphorylation sites are highlighted in red, the BIK1 phosphorylation site in blue. (*B*) Top-view of the BSU1 Kelch domain. Shown is a ribbon diagram colored with a rainbow gradient from blue (N-term) to red (C-term). Kelch motif blades are labeled I to VI, an intrinsically disordered loop is indicated by a dotted line, the BIK1-mediated Ser251 phosphorylation site is shown alongside (in bonds representation, marked by a blue circle). (*C*) Ribbon diagram of the BSU1 phosphatase domain (chain A), with α-helices in blue, β-strands in green, loops in white, and Zn^2+^ ions as gray spheres. The C-terminal tail is shown in dark gray, and Ser764 is depicted in bonds representation, marked by a red circle. ( *D*) Close-up view of the active sites of wild-type BSU1 (left) and the catalytically inactive variant BSU1^421-793^ ^D510N^ ^H707K^ (right). Zn^2+^ ions are shown as green spheres, the distorted trigonal bipyramidal pentavalent coordination is indicated by dotted lines (in magenta), water molecules are shown as red spheres. A phased anomalous difference map calculated from data collected at the Zn K-edge and contoured at 25σ is shown as a green mesh. (*E*) Initial enzyme velocities for BSU1-catalyzed hydrolysis of para-nitrophenyl phosphate (*p*-NPP), plotted as a function of substrate concentration. Data were fitted to the Michaelis-Menten equation by nonlinear regression. Extracted kinetic parameters are summarized in panel *K* (n = 3). (*F*) Metal ion preference of the phosphatase domain using 2 mM *p*-NPP as substrate. Each metal chloride was tested at a fixed concentration of 2 mM. (*G*) BIN2 pTyr dephosphorylation time course experiment using 20 µM BIN2 and 5 µM of either the isolated BSU1 phosphatase domain (top) or the full-length enzyme (bottom). Shown are a immunoblots and coomassie-stained membranes below as loading controls. The presence of BIN2 pTyr200 was detected using a human GSK3 pTyr antibody (n = 2). (*H*) Structural superposition of the BSU1 phosphatase domain (cyan C_α_ trace) and HsPP1γ (in orange, PDB-ID: pdb_00001jk7 (92), r.m.s.d. is ∼0.8 Å comparing 245 corresponding C_α_ atoms). (*I*) Close-up view of the superimposed BSU1 (in cyan) and HsPP1γ (in orange) active sites. Shown are the catalytic Zn^2+^/Mn^2+^ ions (in purple), with coordinating residues depicted in bonds representation. (*J*) Initial enzyme velocities of BSU1 phosphatase domain (or phosphatase-dead) catalyzed hydrolysis of synthetic substrate peptides containing either a pThr (left), pSer (center) or pTyr (right), or of the natural substrate candidates BIN2 and CrCDKB1 (bottom row), using a malachite green-based phosphatase assay (n = 3). (*K*) Table summaries of the enzyme kinetic parameters derived from the experiments shown in *E* and *J* (n.d., no activity detected). (*L*) Metal ion preference of the phosphatase domain vs. synthetic substrate peptides containing either a pThr or pTyr. Assays were performed using 200 µM of peptide substrate and 50 nM of enzyme in the presence of 2 mM of the indicated metal chloride. In the case of ZnCl_2_ + Fe(II)Cl_2_ a final concentration of 500 µM was used (n = 3). (*M*) Close-up view of the RVxF peptide binding grooves in BSU1 (in cyan) and HsPP1α (in orange, PDB-ID: pdb_00003v4y (141), r.m.s.d. is ∼0.8 Å comparing 252 corresponding C_α_ atoms). Shown is a ribbon diagram of the peptide binding groove with substrate peptide-interacting amino-acid side-chains shown in bonds representation. (*N*) Molecular surface view of the BSU1 RVxF peptide binding groove (in cyan) bound to the TEV recognition motif (in yellow, in surface and bonds representation) of a neighboring molecule in the asymmetric unit.

In our structure, the catalytic center of BSU1 coordinates two Zn^2+^ ions (Fig. 1*D*) that co-purified with the enzyme, as previously reported (62). We mutated BSU1 Asp510 to asparagine and His707 to lysine (63) and obtained a 1.6 Å crystal structure of the putatively inactive enzyme (*SI Appendix*, Table S1). The mutations disrupt the coordination of metal ions in the catalytic center without altering the overall structure of the BSU1 core (root mean square deviation, r.m.s.d. is ∼0.7 Å comparing 4,737 corresponding atoms) (Fig. 1*D*). In vitro, BSU1 hydrolyzes the generic phosphatase substrate *p*-Nitrophenyl Phosphate (*p*-NPP) and has a preference for Mn^2+^ ions when assayed against *p*-NPP, as previously reported (34) (Fig. 1 *E* and *F*). The BSU1^421-793^ ^D510N^ ^H707K^ mutant had no detectable activity in this assay (Fig. 1*E*). Next, we purified full-length BIN2, which is auto-phosphorylated on Tyr200 when expressed in *E. coli*, as previously reported (34) (*SI Appendix*, Fig. S2). No dephosphorylation of pTyr200 was observed when BIN2 was incubated with either the isolated BSU1 phosphatase domain, or with full-length BSU1, even at a 4:1 substrate-to-enzyme ratio (Fig. 1*G*).

We therefore characterized the substrate specificity of BSU1 in vitro. Structural homology searches with the program DALI (64) revealed that the BSU1 phosphatase domain most closely resembles protein phosphatase 1 (PP1) enzymes (65, 66), which are generic Ser/Thr phosphatases (67) (Fig. 1*H*) (*SI Appendix* Fig. S3*C*). Notably, the catalytic centers of BSU1 and human PP1 are virtually identical (Fig. 1*I*). Optimal substrate peptides for human PP1 have been previously identified using a high-throughput in vitro dephosphorylation assay (68). Based on this study, we designed three different substrate peptides for BSU1, harboring either a central pSer, pThr or pTyr residue (*SI Appendix*, Fig. S4). We performed enzyme kinetic assays and found that wild-type BSU1 but not the phosphatase-dead BSU1^421-793^ ^D510N^ ^H707K^ mutant efficiently dephosphorylated the pThr-containing peptide, with a catalytic efficiency (k_cat_/K_M_) of ∼24 × 10^3^ M^-1^ s^-1^ (Fig. 1 *J* and *K*). BSU1 also dephosphorylated the pSer-containing peptide (k_cat_/K_M_ ∼4 × 10^3^ M^-1^ s^-1^) but had only very low activity against the pTyr-containing peptide (k_cat_/K_M_ ∼0.05 × 10^3^ M^-1^ s^-1^) (Fig. 1 *J* and *K*). In line with this, the catalytic efficiency for *p*-NPP, a phospho-tyrosine mimicking substrate, is much lower (k_cat_/K_M_ ∼0.09 × 10^3^ M^-1^ s^-1^) than observed for the pThr and pSer-containing substrate peptides (Fig. 1 *E* and *K*). Molecular docking of our synthetic substrate peptides into the Mn^2+^-bound BSU1 phosphatase core using AlphaFold3 (69) revealed that pThr– and pSer-containing peptides may adopt well-defined binding poses within the catalytic center of the BSU1 phosphatase domain, whereas the pTyr-containing peptide was modeled in a distorted conformation to accommodate access to the active site (*SI Appendix*, Fig. S4).

In agreement with the dephosphorylation experiments performed using full-length BIN2 as substrate (Fig. 1*G*), BSU1 showed no detectable activity against a short BIN2 substrate peptide containing pTyr200 (193-KGEANIS(pY)ICSRFYR-207) (Fig. 1 *J* and *K*). Recently, the CYCLIN-DEPENDENT KINASE B1 (CDKB1) was identified as an in vivo substrate for the *Chlamydomonas reinhardtii* Kelch phosphatase BSL1 (58) (*SI Appendix*, Fig. S1). Specifically, Thr14 and neighboring Tyr15 in CrCDKB1 were identified to be hyperphosphorylated in a *bsl1-1* mutant phosophoproteome (58). We found that BSU1 was able to de-phosphorylate a synthetic CrCDKB1 peptide containing pThr14 (k_cat_/K_M_ ∼1 × 10^3^ M^-1^ s^-1^), but not the pTyr15-containing substrate peptide (k_cat_/K_M_ ∼0.04 × 10^3^ M^-1^ s^-1^) (Fig. 1 *J* and *K*). It has been previously reported that the substrate specificity and enzyme activity of animal PP1 are in part determined by the type of metal ions present in the catalytic center (Fig. 1*I*) (70). We therefore assayed BSU1-mediated dephosphorylation of the synthetic pThr– and pTyr-containing substrate peptides in the presence of different metal ions (Fig. 1 *K* and *L*). These assays confirmed BSU1 to represent a Mn^2+^-dependent Ser/Thr protein phosphatase (Fig. 1*L*). Taken together, the BSU1 phosphatase domain shares strong structural homology with human PP1 and behaves as a specific Ser/Thr phosphatase in vitro.

PP1 phosphatases interact with their protein substrates via distinct, conserved binding surfaces including the RVxF motif binding groove (71, 72). BSU1 contains a hydrophobic RVxF binding groove that is highly similar to human PP1 (Fig. 1*M*) and which in our crystal structures is occupied by the C-terminal Tobacco Etch Virus (TEV) protease recognition site that was included in our protein expression construct (Fig. 1*N*). Thus, BSU1 appears to share the catalytic activity and substrate binding mechanism with canonical PP1 enzymes (73).

### Ectopic expression of catalytically inactive BSU1 rescues *bri1-5* mutant phenotypes

Different studies have demonstrated that expression of full-length BSU1 from the *35S* promoter can suppress the growth phenotype of the *bri1-5* BR receptor mutant (33, 34, 36, 39, 40). We used this approach to perform BSU1 structure-function studies, expressing wild-type or mutant fragments of BSU1 fused to a C-terminal 3xFlag tag in the *bri1-5* background (*SI Appendix*, Table S2). We compared the shoot growth phenotypes of the resulting T1 plants to the previously reported *bsu1-1D* allele (36) (Fig. 2*A*). Expression of the isolated BSU1 phosphatase domain (BSU1^421-793^) did partially suppress the *bri1-5* shoot growth phenotype (Fig. 2*A*), while expression of the N-terminal Kelch domain (BSU1^1-357^) further enhanced the stunted growth phenotype of the *bri1-5* mutant (*SI Appendix*, Fig. S5). Surprisingly, the catalytically inactive BSU1 phosphatase domain (BSU1^421-793^ ^D510N^ ^H707K^) (Fig. 1 *E* and *J*) could also suppress the *bri1-5* phenotype (Fig. 2*A*) (*SI Appendix*, Fig. S5). Missense mutations in the RVxF hydrophobic binding groove, such as Cys291 and Leu289 to arginine in human PP1α, have been previously shown to disrupt the interaction of PP1 with protein substrates (74) (Fig. 1*M*). A BSU1 variant, in which we replaced the corresponding Met748 and Ile750 with arginine (BSU1^421-793^ ^M748R^ ^I750R^) (Fig. 1*M*), failed to rescue the *bri1-5* mutant growth phenotype, independent of the catalytic activity of the enzyme (Fig. 2*A*).

**Fig. 2.**
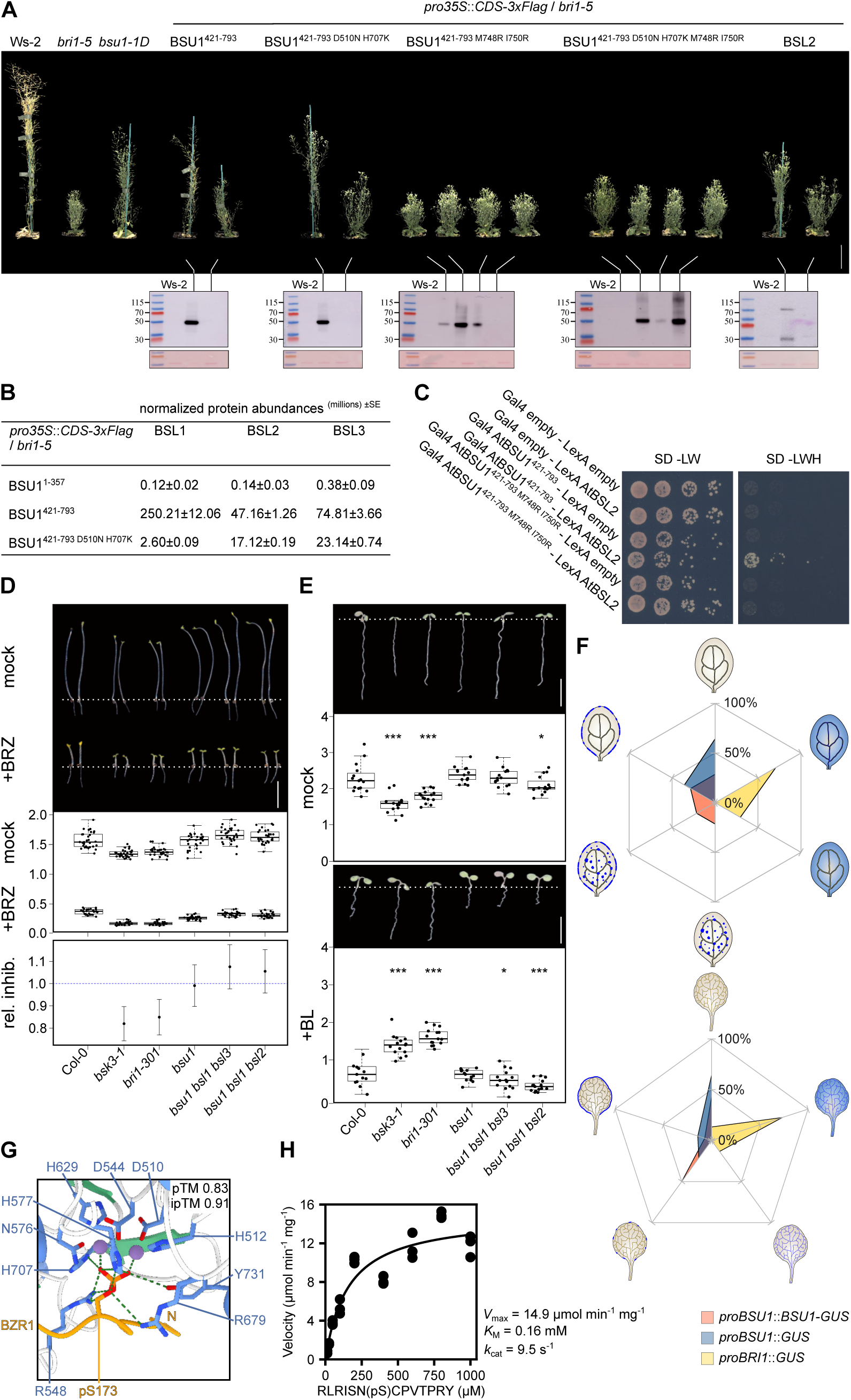
A catalytically inactive BSU1 phosphatase domain can suppress the growth phenotype of the *bri1-5* mutant. (*A*) Shoot growth phenotypes of 45 d-old *bri1-5* plants expressing different BSU1 variants, or full-length BSL2 from the *35S* promoter. The Ws-2 wild type, *bri1-5* (142) and the original *bsu1-1D* (36) allele are shown alongside. BSU1/BSL2 protein expression was assayed by western blot using an α-Flag antibody, the Ponceau-stained membrane is shown below as loading control. (*B*) Relative abundance of BLS1, BSL2 and BSL3 in AP-MS experiments with the Flag-tagged BSU1 Kelch (top), phosphatase (center) or phosphatase-dead (bottom) domains. (*C*) Yeast co-expressing either the wild-type BSU1 phosphatase domain or the mutant carrying point mutations in the RVxF binding groove fused to the Gal4-activation domain (Gal4; prey) and full-length wild-type BSL2 fused to the LexA-binding domain (LexA; bait) were grown on selective SD medium supplemented with histidine (-LW; co-transformation control) or lacking histidine (−LWH; interaction assay). Shown are serial dilutions from left to right after 2d of incubation. (*D*) Hypocotyl growth assay of 5 d-old dark-grown seedlings in the presence and absence of the BR biosynthesis inhibitor BRZ (75). Representative seedlings are shown in the top panel (scale bar = 0.5 cm). Box plots (bold black line, median; box, interquartile range (IQR); whiskers, lowest/highest data point within 1.5 IQR of the lower/upper quartile) are in the center panel with raw data shown as individual points (n = 30). Shown below is relative inhibition of hypocotyl growth in the presence of BRZ plotted together with lower and upper confidence intervals, the dotted horizontal line depicts the Col-0 wild-type reference (rel. inhib., relative inhibition). (*E*) Brassinolide (BL)-mediated root growth inhibition assay. Shown are representative 7 d-old seedlings, grown on ^1/2^MS plates (mock, top panel) or in the presence of 100 nM BL (+BL, bottom panel). Quantifications are shown below as box plots with raw data indicated as individual points (n = 15) and including Dunnett-type (124), two-sided multiplicity-adjusted *p*-values for the comparison against the Col-0 control (\**p* < 0.05; \*\**p* < 0.01; \*\*\**p* < 0.001). (*F*) Spatiotemporal expression patterns of BSU1 and BRI1 reporter lines determined by β-glucuronidase activity (GUS) assays. A sequence ∼1kb upstream of the BSU1 ORF was used to drive *proBSU1*::*BSU1-GUS* and *proBSU1*::*GUS*, respectively. A ∼2kb upstream of BRI1 ORF was used to drive *proBRI1*::*GUS*. 14-17 independent T2 lines for each construct were stained for GUS activity, either using 5 d-old cotyledons (top), or the 4^th^ leaf of 15 d-old soil-grown plants. Shown are distribution star plots with expression patterns divided into categories. (*G*) AlphaFold3-predicted complex structure of BSU1^421-793^ (ribbon diagram) bound to Mn^2+^ ions (purple spheres) and BZR1^168-178^pS (in orange). Predicted salt bridges between the phosphate group and BSU1 residues are depicted as dotted lines (in green). AlphaFold3 pTM and ipTM values are reported alongside. (*H*) Initial enzyme velocities of BSU1 phosphatase domain catalyzed hydrolysis of the synthetic BZR1^168-178^pS substrate peptide and including enzyme kinetic parameters (n = 3).

Our observation that the presence of an intact substrate binding groove allows the BSU1 phosphatase domain to rescue the *bri1-5* phenotype prompted us to investigate if known BR signaling components may interact with BSU1 when over-expressed in the *bri1-5* mutant. We performed affinity purification followed by mass spectrometry (AP-MS) experiments with the N-terminal Kelch domain, the C-terminal phosphatase domain, and the catalytically inactive BSU1^421-793^ ^D510N^ ^H707K^, respectively. With the exception of the BSU1 homologs BSL1-3, no known BR signaling components were recovered. The BSL1-3 proteins were strongly enriched in the BSU1 phosphatase and phosphatase-dead samples, but not in the Kelch domain sample (Fig. 2*B*). Full-length BSU1 has been previously shown to interact with BSL1-3 in vitro and in planta (40). The interaction surface on BSU1 has not yet been mapped. We found that the wild-type BSU1 phosphatase domain, but not the BSU1^421-793^ ^M748R^ ^I750R^ mutant targeting the RVxF binding groove interacted with BSL2 in yeast two-hybrid assays (Fig. 2*C*). Together, these experiments suggest that the over-expression of BSU1 suppresses the phenotypes of the BR receptor mutant allele *bri1-5* through RVxF binding groove-mediated protein-protein interactions rather than via its catalytic phosphatase activity (Fig. 2*A-C*).

The presence of BSLs in our AP-MS assays led us to reanalyze their role in BR signaling in relation to BSU1. We used CRISPR/Cas9 gene editing in Col-0 background to generate a *bsu1* knock-out allele, and higher-order *bsu1 bsl* mutants (*SI Appendix*, Fig. S6). *bsu1* mutants were indistinguishable from wild-type plants and exhibited normal BR responses (Fig. 2 *D* and *E*), as previously described (34, 36). *bsu1 bsl1 bsl3* and *bsu1 bsl1 bsl2* triple mutants had significantly more stomata compared to wild type (*SI Appendix*, Fig. S7), in good agreement with the stomatal patterning phenotype previously reported for a *bsl1 bsl2 bsl3* T-DNA insertion mutant (52). *bsu1 bsl1 bsl3* and *bsu1 bsl1 bsl2* mutants showed no significant differences from the wild-type control in quantitative hypocotyl growth assays performed in the presence and absence of the BR biosynthesis inhibitor brassinazole (BRZ) (75, 10, 15, 19, 24) (Fig. 2*D*). Addition of the potent brassinosteroid brassinolide (BL) inhibited primary root growth of *bsu1, bsu1 bsl1 bsl3* and *bsu1 bsl1 bsl2* mutant plants to wild type-like levels, while the known BR signaling mutants *bsk3-1* (30) and *bri1-301* (76) were insensitive (Fig. 2*E*). Both *bsu1 bsl1 bsl3* and *bsu1 bsl1 bsl2* mutants had shorter siliques compared to wild type (*SI Appendix*, Fig. S8*A*), indicative of a fertility defect, as previously described (37). We generated *bsu1 bsl1 bsl3 bsl2*^+/-^ or *bsu1 bsl1 bsl2 bsl3*^+/-^ plants, but could not isolate a homozygous quadruple mutant. In all higher-order *bsu1 bsl* mutants we observed a severe seed-abortion phenotype, suggesting that the quadruple mutant is likely nonviable and therefore cannot be evaluated for BR responses (*SI Appendix*, Fig. S8 *B* and *C*). Overall, our experiments provide no genetic evidence for a function of BSU1 and BSL protein phosphatases as positive regulators of BR signaling, while confirming their role in stomatal patterning and in other, essential cell signaling processes.

Different experimental approaches have defined the BR receptor BRI1 to be an ubiquitously expressed receptor kinase throughout plant development (77–81). However, there are conflicting reports regarding the tissue expression of *BSU1* (36, 37). We generated *proBSU1*::*GUS* and *proBSU1*::*BSU1-GUS* reporter lines and found *BSU1* expression to be low and restricted to the outer leaf margin in cotyledons and true leaves across ∼15 independent T2 lines (Fig. 2*F*) (*SI Appendix*, Fig. S9). In contrast, a *proBRI1*::*GUS* reporter line confirmed the ubiquitous expression of *BRI1* (Fig. 2*F*) (*SI Appendix*, Fig. S9). These experiments indicate that the expression patterns of the BR receptor and its previously proposed downstream signaling component BSU1 are largely non-overlapping.

In conclusion, the expression domains of BSU1 and BRI1 only partially overlap (Fig. 2*F*) and *bsu1* and *bsu1 bsl* loss-of-function mutants display wild type-like BR responses (Fig. 2 *D* and *E*). Over-expression of phosphatase-dead BSU1 can rescue the growth phenotypes of the BR receptor allele *bri1-5* (Fig. 2*A*). When over-expressed, BSU1 can interact with BSL1-3, as previously reported (40), and this interaction and the ability to suppress *bri1-5* requires a functional RVxF binding groove (Fig. 2 *B*, *C* and *F*). Thus, the interaction between ectopically expressed BSU1 and BSLs may constitute a key molecular step in reestablishing intracellular BR signaling in the *bri1-5* mutant. In line with this, over-expression of BSL2 can likewise partially suppress the *bri1-5* mutant growth phenotype (Fig. 2*A*). It is thus possible, that the *bsu1-D* allele induces the artificial over-accumulation of Kelch Ser/Thr phosphatase activity (Fig. 1), which in turn could lead to a direct dephosphorylation of pSer/pThr containing BR signaling components. We attempted to identify potential pSer or pThr-containing substrates, by docking known BR signaling components (2) to the BSU1 phosphatase domain using AlphaFold3 (69). The BR transcription factor BZR1 was identified as a high scoring interaction, with an interface predicted Template Modeling (ipTM) score of ∼0.9 (Fig. 2*G*). Notably, the previously characterized BZR1^168-178^pS peptide (24), which contains a GSK3 phosphorylation site at Ser173, and which has been previously shown to be phosphorylated by BIN2 (82, 83), resembles a bona fide enzyme – substrate complex when docked to the BSU1 phosphatase domain (Fig. 2*G*). In accordance, BSU1 very efficiently dephosphorylated BZR1^168-178^pS in vitro, with a k_cat_/K_M_ of ∼59 × 10^3^ M^-1^ s^-1^. (Fig. 2*H*). BSL1-3 could not be evaluated in this assay because we were unable to purify these BSU1 homologs (which share ∼65% sequence identity in their phosphatase domains) from either *E. coli* or insect cells, as previously reported for other PP1 enzymes (73). Taken together, artificial over-accumulation of BSU1 and BSLs may result in the direct dephosphorylation of pBZR1/pBES1 in the *bsu1-D* mutant, as initially proposed (36).

### A PP1-like regulatory tail in BSU1 is phosphorylated by the CDK1 – cyclin B1 – CKS1 complex

Analysis of the crystal structure of the wild-type BSU1 phosphatase domain revealed a well-ordered C-terminal tail that folds back into the active site of the enzyme (Figs. 1*A* and 3*A*). BSU1 Thr785 is phosphorylated, and its phosphate moiety is coordinated by Arg548, Arg679 and Tyr731 (Fig. 3*A*), which are conserved among BSU1, BSLs and human PP1 isoforms (*SI Appendix*, Fig. S1). Despite its proximity to the catalytic metal center, pThr785 is too far from both the metal-activated water molecule that mediates the nucleophilic attack of PP1 substrates (66) and the general base His577 (which corresponds to His125 in HsPP1α) to enable catalysis (Fig. 3*A*). Mass spectrometric analysis of insect cell and plant extracted BSU1 revealed that Thr785 is also phosphorylated in vivo (Fig. 3*B*) (*SI Appendix*, Table S3). Notably, BSU1 and PP1 isoforms have very similar C-terminal tails, with BSU1 pThr785 mapping to the previously identified PxTPP motif in PP1 (Fig. 3*C*) (84). Phosphorylation of the corresponding Thr320 in HsPP1α is an important control mechanism for cell-cycle progression, for example regulating the phosphorylation of the retinoblastoma protein during G1 to S phase transition (84). It has been previously demonstrated that the cyclin-dependent kinase 1 (CDK1) – cyclin B1 complex phosphorylates the conserved threonine residue within the PxTPP motif in human and fission yeast PP1 (85–87). Consistent with this, a recombinantly expressed and purified CDK1 – cyclin B1 – cyclin-dependent kinase subunit 1 (CKS1) complex (88) readily phosphorylated the isolated BSU1 phosphatase domain, but not the BSU1^421-793^ ^T785A^ mutant protein (Fig. 3*D*).

**Fig. 3.**
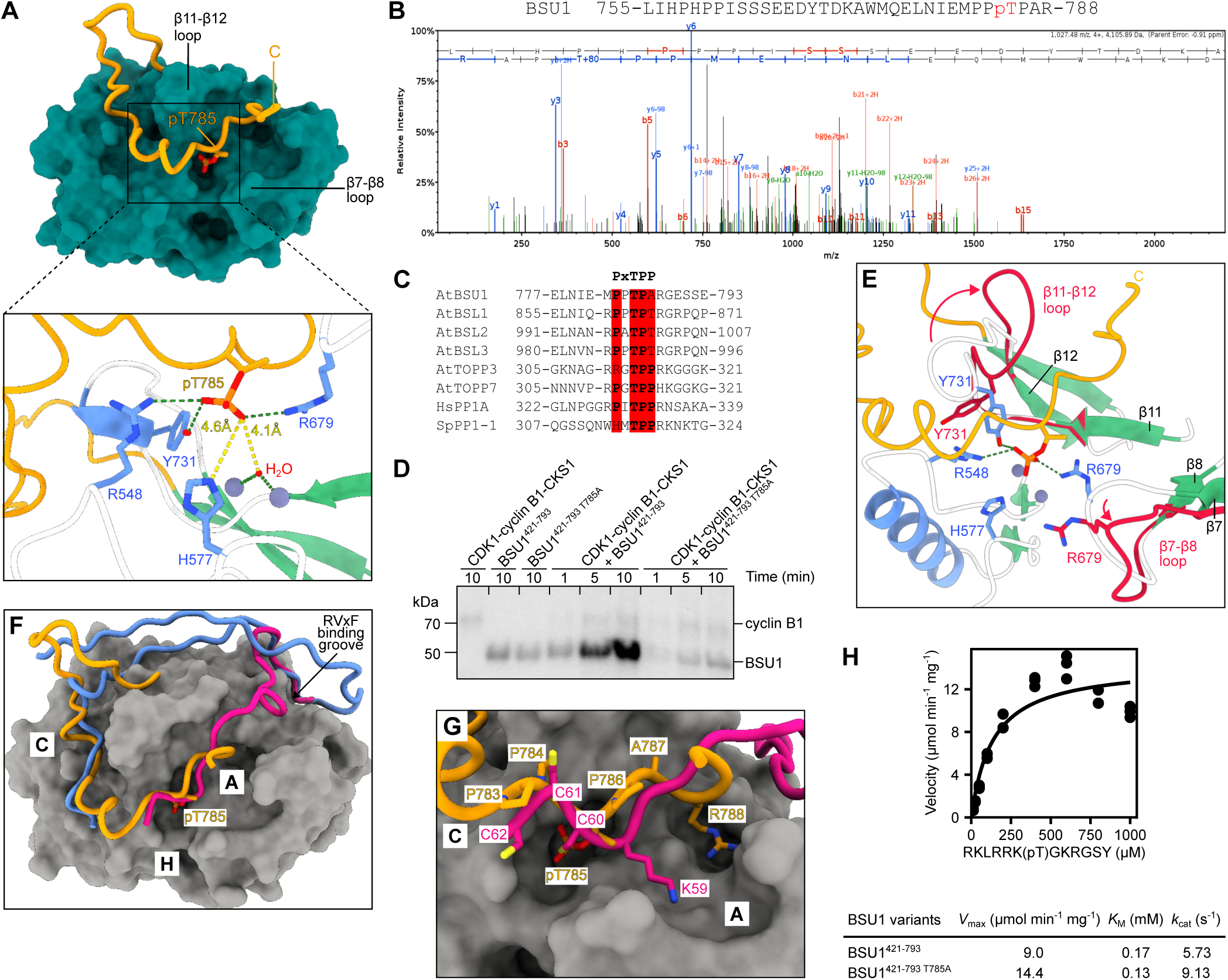
The C-terminal tail of BSU1 is phoshorylated by the CDK1 – cyclin B1 – CKS1 complex. (*A*) Molecular surface view of the BSU1 phosphatase domain (residues 450-755), the C-terminal tail (residues 756-793) is shown as a ribbon diagram (in orange), pThr785 is inserted into the active site (in bonds representation). The relative positions of two substrate-binding loops are indicated. The inset provides a close-up view of the BSU1 active site with two divalent metal ions and the catalytic water molecule shown as purple and red spheres, respectively. Residues contacting pThr785 (in orange) are highlighted in bonds representation (in blue), hydrogen bonds are shown as green dotted lines. The closest distances between the phosphate moiety of pThr785 and the catalytic water molecule or the general base His577 are indicated by yellow dotted lines. (*B*) MS1 spectrum of the phosphorylated BSU1^755-788^ peptide. (*C*) Multiple sequence alignment of the C-terminal tail of BSU1, BSL1-3, two PP1-like TOPP phosphatases from *Arabidopsis thaliana*, and human and fission yeast PP1. The position of the PxTPP motif containing the pThr site is indicated (in red). (*D*) Autoradiograph of an in vitro transphosphorylation time course assay. 10 nM of the human CDK1 – cyclin B1 – CKS1 complex were incubated with 10 µM BSU1^421-793^ or with BSU1^421-793^ ^T785A^, where Thr785 in the PxTPP motif was replaced by an alanine, at 37 °C for the indicated time. ( *E*) Structural superposition of BSU1^421-793^ (ribbon diagram with α-helices shown in blue, β-strands in green and loops in white) and BSU1^421-793^ ^D510N^ ^H707K^ (in red, r.m.s.d. is ∼0.4 Å and 2.7 Å comparing 295 and 311 corresponding C_α_ atoms, respectively). Loop regions undergoing conformational changes (in red) upon binding of the phosphorylated C-terminal tail in BSU1^421-793^ (in orange) are labeled and indicated by red arrows. (*F*) Structural superposition of BSU1^421-793^ phosphatase domain (residues 450-755, gray molecular surface; residues 756-793, orange ribbon diagram), the HsPP1α – spinophilin complex (in blue, spinophilin residues 425-474, r.m.s.d. is ∼0.8 Å comparing 228 corresponding C_α_ atoms, PDB-ID: pdb_00003egg) (94), and the HsPP1α – Inhibitor-3 complex (in magenta, Inhibitor-3 residues 39-62, r.m.s.d. is ∼0.8 Å comparing 245 corresponding C_α_ atoms, PDB-ID: pdb_00008dwk) (62). The positions of the C-terminal (**C**), acidic (**A**) and hydrophobic (**H**) binding grooves in PP1 enzymes are indicated (73). (*G*) Close-up view of the BSU1 substrate binding pocket (gray molecular surface) shown together with the phosphorylated C-terminal tail (in orange) and the C-terminus of Inhibitor-3 (in magenta, residues 59-62 shown in bonds representation, PDB-ID: pdb_00008dwk) (62). (*H*) Initial enzyme velocities of the BSU1 phosphatase domain or of the BSU1 Thr785Ala mutant vs a synthetic pThr-containing substrate peptide, and including the derived enzyme kinetic parameters (n = 3).

Mutation of Thr320 in the C-terminal tail to alanine constitutively activates human PP1α (89) and phosphorylation of this threonine residue correlates with a reduction of PP1 activity (87, 90). However, no structure of a PP1 with an ordered C-terminal tail has been resolved thus far (91). Our structure of the active BSU1 phosphatase domain now reveals that the phosphorylated C-terminus adopts an inhibitor-like binding mode, triggering rearrangement of the β7-β8 and β11-β12 loops, as previously described for small molecule and protein inhibitors (92, 93) (Fig. 3 *A* and *E*). We therefore compared BSU1 with previously resolved PP1 – inhibitor complex structures, identified with DALI (64) (*SI Appendix*, Fig. S3). We found that part of the BSU1 C-terminal tail (residues 767-778) binds the PP1 C-terminal binding groove, which is also targeted by, for example, the neuronal PP1 inhibitor protein spinophilin (94) (Fig. 3*F*). In addition, the phosphorylated BSU1 C-terminal tail occupies the acidic binding groove, adopting a similar conformation as recently reported for human Inhibitor-3 (62). (Fig. 3*G*). Both the triple Cys-motif of Inhibitor-3 and the ^783^PPpTP^786^ segment of BSU1 bind across the active site of the enzyme, while two basic residues (Arg788 in BSU1 and Lys59 in Inhibitor-3) target the acidic binding groove (Fig. 3*G*). Spinophilin and Inhibitor-3 have been shown to inhibit the activity of human PP1α by blocking substrate binding grooves and access to the active site (62, 94). By structural analogy, this implies that phosphorylation of Thr785 in the C-terminal tail of BSU1 inhibits both the binding of protein substrates, and their dephosphorylation. Consequently, the BSU1^421-793^ ^T785A^ mutant dephosphorylated a synthetic pThr-containing substrate peptide about two times more efficiently than the BSU1^421-793^ wild-type phosphatase domain, with k_cat_/K_M_ values of ∼70 × 10^3^ M^-1^ s^-1^ and ∼35 × 10^3^ M^-1^ s^-1^, respectively (Fig. 3 *H* and 1*K*).

Together, the structural homology of BSU1 to PP1 enzymes, its phosphorylation by the CDK1 – cyclin B1 – CKS1 complex, and the presence of conserved substrate-binding and regulatory motifs together support a role for plant Kelch phosphatases in cell cycle regulation. In line with this, we observed induction of BSU1 expression and protein accumulation in callus, a tissue formed by rapidly growing and dividing cells (*SI Appendix*, Fig. S10).

### The Kelch phosphatase BSLM regulates the Marchantia cell cycle

Our observation that Arabidopsis *bsu1 bsl1 bsl2 bsl3* quadruple mutants are not viable (*SI Appendix*, Fig. S8) prompted us to study the function of Kelch phosphatases in *Marchantia polymorpha.* Marchantia Kelch phosphatases are encoded on the sex chromosome and we named them MpBSLM (male, MpVg00440 https://marchantia.info) and MpBSMF (female, MpUg00300), respectively. During the gametophyte phase MpBSLM appears as a single copy gene in the Tak-1 wild type. *Marchantia polymorpha* lacks a canonical BR receptor and does not respond to brassinosteroids (95). However, Marchantia harbors a BIN2-like GSK3 kinase, MpGSK, and a MpBES1 transcription factor (95, 96). Mp*gsk*^ge^ and Mp*bes1*^ge^ knock-out lines are both strongly affected in growth and cell differentiation (95, 96).

We generated Mp*bslm*^ge^ and Mp*gsk*^ge^ knock-out mutants using CRISPR/Cas9 gene editing and an amiRNA-MpBSLM knock-down line (*SI Appendix*, Fig. S11). Mp*bslm*^ge^ and Mp*gsk-4*^ge^ mutants both had an undifferentiated cell mass phenotype (Fig. 4*A*), similar to what has been previously described for the Mp*gsk-1*^ge^ and Mp*gsk-2*^ge^ alleles (96). The amiRNA-MpBSLM mutant displayed an intermediate growth phenotype (Fig. 4*A*). The Mp*bslm*^ge^ and Mp*gsk-3*^ge^ mutants had no recognizable air pores on the thallus surface (Fig. 4*B*). The typical thallus body plan with a dorsal epidermal layer, a central storage layer and a ventral epidermis could not be differentiated in either the Mp*bslm*^ge^ or the Mp*gsk-3*^ge^ mutant, respectively (Fig. 4*B*). The ventral and dorsal epidermis and some air chambers were still recognizable in the weaker amiRNA-MpBSLM allele (Fig. 4*B*).

**Fig. 4.**
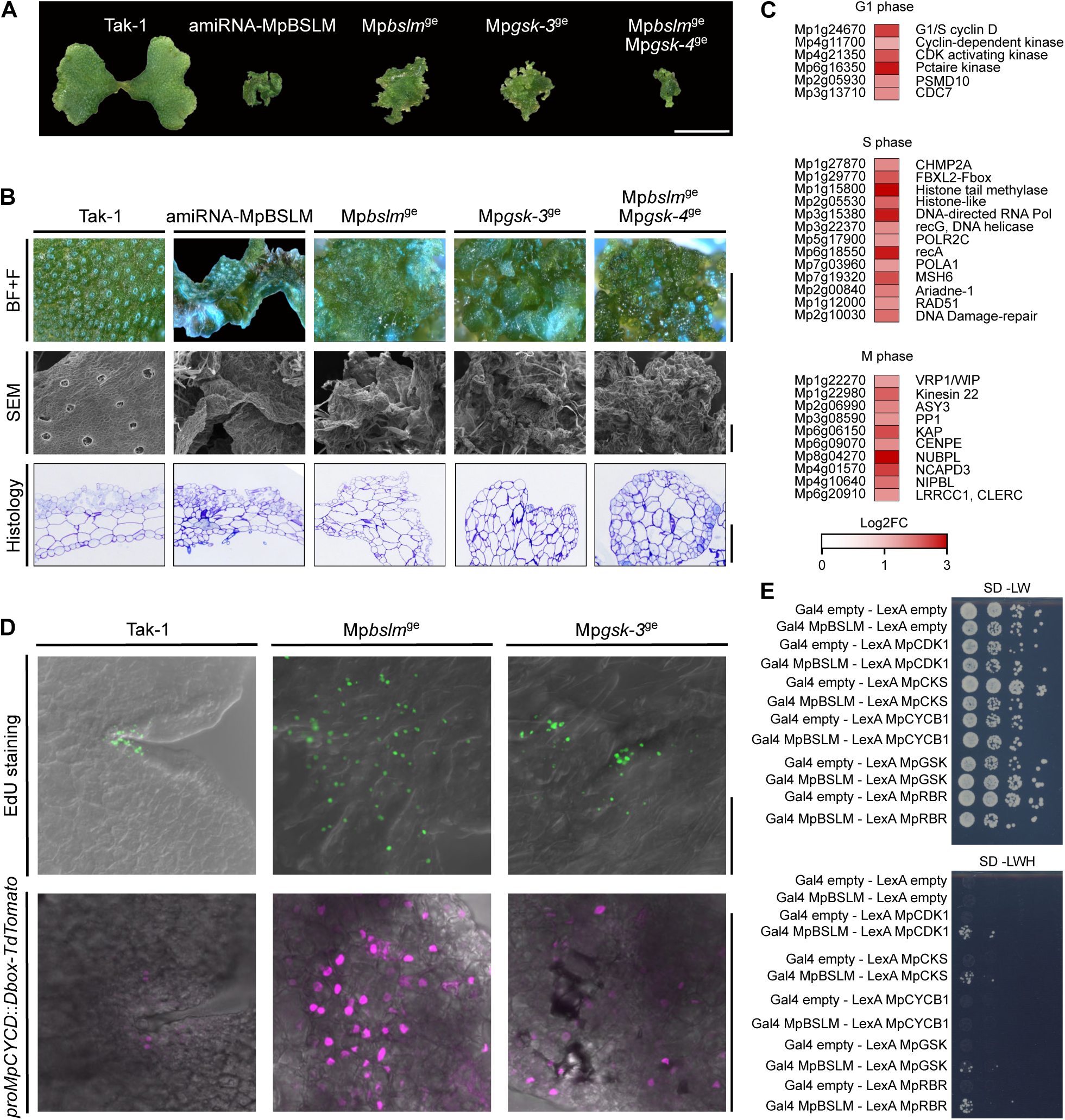
The Kelch phosphatase BSLM is a cell cycle regulator in Marchantia. (*A*) Representative thallus phenotypes of 14 d-old Tak-1 and amiRNA-MpBSLM plants grown from gemma, and Mp*bslm*^ge^, Mp*gsk-3*^ge^ and Mp*bslm*^ge^ Mp*gsk-4*^ge^ propagated from thallus stubs (scale bar = 1 cm). (*B*) Ultrastructural phenotypes of 14 d-old plants. On top, Bright Field + Fluorescence (BF+F) thallus views showing the air pore distribution across the genotypes from A (pore autofluorescence in blue, scale bar = 500 µm). Center: Scanning Electron Microscopy (SEM) views of selected thallus areas (scale bar = 200 µm). Bottom: Technovit7100-fixed histology thallus cross sections stained with toluidine blue (scale bar = 10 µm). (*C*) Heat map of selected up regulated genes putatively involved in cell cycle regulation, identified by RNAseq comparing 14 d-old Tak-1 and Mp*bslm*^ge^, log_2_FC > 1 and FDR < 0.01. (*D*) Top: Combined bright-field view (in gray scale) and EdU staining (in green) of 3 d-old thalli (scale bar = 200µm). Bottom: Expression of the *proMpCYCD*::*Dbox-TdTomato* cell cycle reporter (in magenta) in Tak-1, and in Mp*bslm*^ge^ or Mp*gsk-3*^ge^ double transgenic lines (scale bar = 50 µm). (*E*) Yeast co-expressing MpBSLM fused to the Gal4-activation domain (Gal4; prey) and MpCDK1 (Mp2g04270), MpCKS (Mp1g16700), MpCYCB1 (Mp5g10030), MpGSK (Mp7g04170) or MpRBR (Mp8g18830) fused to the LexA-binding domain (LexA; bait) were grown on selective SD medium supplemented with histidine (-LW; co-transformation control) or lacking histidine (−LWH; interaction assay). Shown are serial dilutions from left to right after 3 d of incubation.

Arabidopsis has four Kelch phosphatases and 10 BIN2-like GSK3/SHAGGY kinases (36, 97). The presence of a single *BSL* and *GSK* gene in Tak-1 and their highly similar loss-of-function phenotypes prompted us to investigate their genetic interaction. We generated a Mp*bslm*^ge^ Mp*gsk-4*^ge^ double mutant, which displayed a more severe growth defect compared to either the Mp*bslm*^ge^ or the Mp*gsk-3*^ge^ single mutant, respectively (Fig. 4 *A* and *B*).

We next performed RNA-seq analyses comparing Mp*bslm*^ge^ to the Tak-1 wild-type. We found ∼500 and ∼800 genes to be significantly up– or down-regulated, respectively (*SI Appendix*, Fig. S12 and Table S4). Among the differentially expressed genes, we identified several cell cycle components and cell-cycle regulated genes, including a Marchantia cyclin-dependent kinase and a cyclin D isoform (Fig. 4*C*).

To investigate if the Mp*bslm*^ge^ and Mp*gsk-3*^ge^ mutants were enriched with cells undergoing the cell cycle we first used 5-Ethynyl-2’-deoxyuridine (EdU) staining to mark cells undergoing DNA duplication during a 3 h time period, as previously described (98). We found EdU staining to be restricted to the meristem of Tak-1 (Fig. 4*D*) (98). In both the Mp*bslm*^ge^ and the Mp*gsk-3*^ge^ mutant we observed EdU stained cells across the entire thallus surface and no distinct meristematic region could be identified (Fig. 4*D*). A technical limitation of EdU staining is that it captures DNA synthesis only during the labeling window and remains detectable in cells even after DNA replication has been completed. To overcome these issues, we used the previously reported *proMpCYCD*::*Dbox-TdTomato* cell cycle marker (96), which is specifically active during the G1-S phase. We transformed the reporter construct into Tak-1 and our Mp*bslm*^ge^ and Mp*gsk-3*^ge^ mutants. We observed only a weak TdTomato signal in the Tak-1 wild-type, which was restricted to the meristem (Fig. 4*D*). In contrast, a much stronger TdTomato signal was observed in both the Mp*bslm*^ge^ and Mp*gsk-3*^ge^ mutants (Fig. 4*D*). In line with the EdU staining results, also the *proMpCYCD*::*Dbox-TdTomato* reporter was expressed throughout the thallus (Fig. 4*D*). In agreement with our biochemical analysis of BSU1 in Arabidopsis (Fig. 3*D*), MpBSLM weakly interacted with MpCDK1 and MpCKS, but not with MpCYCB1 in yeast two-hybrid assays (Fig. 4E). Additionally, we found MpBSLM to weakly interact with the Marchantia retinoblastoma protein MpRBR (Fig. 4*E*), a known PP1 substrate in human cells (84). Together, these experiments suggest a function for MpBSLM in Marchantia cell cycle regulation. Deletion of *MpBSLM* results in cell over-proliferation and uncontrolled cell differentiation. The Mp*gsk-3*^ge^ allele has an overall similar phenotype (Fig. 4*A*) and MpBSLM and MpGSK weakly interact in yeast two-hybrid assays (Fig. 4*E*). However, the enhanced growth phenotype of the Mp*bslm*^ge^ Mp*gsk-4*^ge^ double mutant implies that MpBSLM and MpGSK may not function in a strictly linear signaling pathway (Fig. 4*A*).

## Discussion

The *Arabidopsis thaliana* Kelch phosphatase BSU1 was initially characterized as an okadaic acid-sensitive, PP1-like Ser/Thr phosphatase (36, 92). Later work positioned BSU1 as a key component of the intracellular brassinosteroid (BR) signaling pathway. Specifically, BSU1 was proposed to dephosphorylate the pTyr200 residue of BIN2, thereby regulating its activity toward the transcription factors BZR1 and BES1 (33, 34). However, based on our structural and enzyme kinetic analyses, the BSU1 phosphatase domain consistently behaves as PP1-like Ser/Thr phosphatase and we could not detect dephosphorylation of BIN2 pTyr200, either in the full-length kinase or in a synthetic peptide containing this site (Fig. 1*G*, *J–L*).

It has been previously demonstrated that constitutive over-expression of full-length wild-type BSU1 can partially suppress the growth phenotypes of both weak and strong BR receptor alleles (33, 34, 36, 40). Our structure-function experiments now reveal that (i) expression of the isolated Kelch domain further exacerbates the growth phenotype of the *bri1-5* mutant (*SI Appendix*, Fig. S5), (ii) the isolated BSU1 phosphatase domain alone is sufficient to suppress this growth defect (Fig. 2 *A*) and (iii) a catalytically inactive BSU1 variant retains the ability to suppress the *bri1-5* phenotype (Figs. 1*E* and 2*A*). Together, these results, along with our observations that *bsu1* knockout alleles have no detectable BR signaling defects and that *BSU1* is not broadly expressed, are inconsistent with a central role for BSU1 in canonical BR signal transduction (2, 34). Instead, they support the interpretation that *bsu1-1D* and *pro35S*::*BSU1* act as neomorphic alleles in this context. This raises the question of how over-expressing a catalytically inactive phosphatase core can partially restore BR signaling in the *bri1-5* allele (Fig. 2*A*). Our findings that (i) a BSU1 variant with a disrupted RVxF substrate-binding groove fails to suppress the *bri1-5* phenotype (Figs. 1*M* and 2*A*), (ii) BSL1, BSL2 and BSL3 associate with the overexpressed BSU1 phosphatase domain in *bri1-5* plants (Fig. 2*B*), and (iii) the RVxF binding groove mediates the interaction between BSU1 and BSL2 (Fig. 2*C*), suggest that BSU1 overexpression in *bri1-5* may influence BSL accumulation, distribution, or sub-cellular localization, as previously suggested (40). This, in turn, could indirectly alter BR signaling. In line with this, over-expression of BSL2 also suppresses the *bri1-5* phenotype (Fig. 2*A*). However, in agreement with an earlier report (37), our newly generated *bsu1 bsl1 bsl3* and *bsu1 bsl1 bsl2* knock-out mutants had no detectable BR phenotypes, even in highly sensitive and quantitative hypocotyl growth assays (Fig. 2*D*), which we have previously used to detect very weak BR signaling defects in accessory BR signaling components such as BIR receptor pseudokinases or 14-3-3 proteins (10, 24). Importantly, the triple mutants display the previously reported stomatal patterning defects (52) (SI Appendix, Fig. S7). The fertility defect of *bsu1 bsl1 bsl3 bsl2*^+/-^ and *bsu1 bsl2 bsl3*^+/-^ mutants limits our genetic analysis of BSL isoforms in BR signaling (*SI Appendix*, Fig. S8). It remains formally possible that a single *BSL2* or *BSL3* allele can support BR signaling to wild type-like levels (Fig. 2 *D* and *E*).

While our loss-of-function studies cannot substantiate a role for BSU1 or BSLs in BR signaling, over-expression of these phosphatases can clearly impact the BR pathway (Fig. 2A and *SI Appendix*, Fig. S5). One possible mechanism would be that BSU1 and BSL overaccumulation leads to an increased dephosphorylation of BR signaling components containing pSer and/or pThr sites. We found that the BSU1 phosphatase domain can efficiently dephosphorylate pBZR^169-182^ in vitro, a fragment containing an important BIN2 phosphorylation site in BZR1 required for 14-3-3 binding and nuclear-cytoplasmic shuttling (24, 45, 82, 83, 99). Dephosphorylation of the related BES1 transcription factor by BSU1 in vivo and in vitro has been previously reported (36), and accumulation of dephosphorylated BES1/BZR could rationalize the suppression of the *bri1-5* mutant phenotype in BSU1 or BSL2 over-expressing lines. An alternative possibility is that a BR-regulated protein kinase preferentially phosphorylates BSU1, BSL2, or BSU1-BSL complexes in our *bri1-5* rescue lines (Fig. 2*A*) (40), protein assemblies that would not normally be present in cells or tissues actively engaged in BR signaling (Fig. 2*F*). Indeed, the recombinant BIN2 kinase can efficiently trans-phosphorylate BSU1, especially its N-terminal Kelch domain (*SI Appendix*, Fig. S13). Taken together, several molecular scenarios could account for the enhanced BR signaling observed in transgenic lines over-accumulating Kelch phosphatases. However, our genetic experiments and previous studies (37) do not provide compelling evidence for the involvement of these broadly acting Ser/Thr phosphatases in BR signaling.

While the molecular functions of Arabidopsis BSU1/BSLs in cell signaling (52, 54, 55) warrant further investigation, a core function for Kelch phosphatases in cell cycle regulation has been established in Chlamydomonas (57, 58). Specifically, CrBSL1 (*SI Appendix*, Fig. S1) has been reported to promote mitotic entry by dephosphorylating the cyclin-dependent kinase CDKB1 (58). We found that the BSU1 phosphatase core is structurally highly similar to human and yeast PP1, well-established cell cycle regulators that counteract CDK1-mediated phosphorylation events (89, 90). The HsPP1α and BSU1 active sites are virtually identical and BSU1 harbors the known PP1 RVxF, C-terminal, acidic and hydrophobic binding grooves (Figs. 1 *H* and *I*, 3*F*). While the structure of an apo PP1 enzyme has not been resolved so far (94), our structure of a catalytically inactive BSU1 reveals the substrate-binding loops in an open conformation (Fig. 3*E*).

In the crystal structure of the active BSU1 phosphatase domain, we detected phosphorylation of the regulatory C-terminal tail at the previously identified PxTPP motif (84), with the tail occupying the active site and interacting with both the C-terminal and acidic binding grooves (Fig. 3 *A-C* and *F*). The phosphorylated C-terminal tail can block both access to the PP1 active site and restrict the binding of selected substrates and inhibitors (Fig. 3 *F* and *G*). Mutation of the phosphorylated BSU1 Thr785 to alanine increased the dephosphorylation of a pThr-containing substrate peptide (Fig. 3*H*), rationalizing previous genetic and biochemical studies (84, 87, 89, 90). Our structural data further support a recent mechanistic analysis of a PP1 C-terminal tail deletion mutant (100).

The central threonine residue in the PxTPP motif has been previously shown to be phosphorylated by CDK1 (85, 86), with CDK1 counteracting PP1 activity (101). Consequently, BSU1 Thr785 was efficiently phosphorylated by the CDK1 – cyclin B1 – CKS1 complex in vitro (Fig. 3 *D*). While we originally identified the pThr785 site in insect cell-expressed BSU1, this site is also found phosphorylated in Arabidopsis (*SI Appendix*, Table S3), and in CrBSL1 in Chlamydomonas (58), suggesting that the PP1-like inhibitory mechanism is conserved among plant Kelch phosphatases.

Our structural and biochemical data strongly argue for a function of Kelch phosphatases as cell cycle regulators, similar to PP1 in yeast and human. Consistent with this notion, knockout of the single Kelch phosphatase gene in *Marchantia polymorpha* Tak-1 results in a severe phenotype, with the Mp*bslm*^ge^ mutant forming a drastically undifferentiated cell mass (Fig. 4 *A* and *B*). In agreement with the recent study in Chlamydomonas (58), our work supports a core function for Kelch phosphatases in algal and plant cell cycle progression, likely via multiple regulatory mechanisms. Whether Kelch phosphatases share redundant functions with type-one serine/threonine protein phosphatases (TOPPs) in plants remains to be characterized (102).

## Materials and methods

### Expression construct cloning

The Kelch domain (residues 1–357; BSU1^1-357^) and the full-length enzyme (residues 1–793; BSU1^1-793^) from *Arabidopsis thaliana* BSU1 (TAIR-ID: At1g03445, https://www.arabidopsis.org) were cloned from codon-optimized synthetic genes (Twist Bioscience) into the pBB3 vector for expression in *Spodoptera frugiperda* Sf9 cells. The constructs carried an N-terminal 10×His-Twin-StrepII affinity tag followed by a tobacco etch virus protease (TEV)-cleavable site. The phosphatase domain (residues 421–793; BSU1^421-793^) was cloned into the pBB40 vector carrying a C-terminal TEV-cleavable Twin-StrepII-10×His affinity tag (*SI Appendix*, Table S2).

The full-length BIN2 kinase (residues 1–380; BIN2) from *Arabidopsis thaliana* was cloned into the pMH_HStrx vector from a synthetic gene codon-optimised for expression in *Escherichia coli*. The vector provides an N-terminal 10×His-Streptavidin II-thioredoxin tag, followed by a TEV-cleavage site. Point mutations in BSU1^421-793^ and BIN2 were generated by site-directed mutagenesis and subcloned into plasmids pBB40 and pMH_HStrx, respectively. The phosphatase-dead BSU1^421-793^ variant was produced by replacing the Asp510 and His707 residues in the catalytic center with asparagine and lysine, respectively (BSU1^421-793^ ^D510N^ ^H707K^). The kinase-dead BIN2 variant was generated by replacing the catalytic Lys69 with arginine (BIN2^K69R^), and the non-phosphorylatable BIN2 mutant was obtained by replacing Tyr200 with phenylalanine (BIN2^Y200F^).

The synthetic genes encoding full-length CYCLIN-DEPENDENT KINASE-1 ISOFORM 1 (residues 1–297; HsCDK1), G2/MITOTIC-SPECIFIC CYCLIN-B1 (residues 1–433; HsCyclin B1) and CYCLIN-DEPENDENT KINASES REGULATORY SUBUNIT-1 (residues 1–79; HsCKS1) from *Homo sapiens* were obtained as gene optimized versions for expression in Sf9 cells (Thermo Fisher Scientific) as previously described (88). HsCDK1, HsCyclin B1 and HsCKS1 were cloned into the pF1 vector for co-expression forming the CDK1-cyclin B1-CKS1 complex. An 8×His tag was fused to the C-terminus of HsCDK1 and a TEV-cleavable Twin-StrepII tag was fused to the C-terminus of HsCyclin B1.

### Protein expression and purification

BSU1^1-357^, BSU1^421-793^, BSU1^421-793^ ^D510N^ ^H707K^, and BSU1^1-793^ were expressed in *Spodoptera frugiperda* Sf9 cells using recombinant baculoviruses. Typically, 10 mL of recombinant P3 baculovirus was used to infect 200 mL of Sf9 insect cells growing in SF900 media (Gibco) at a cell density of ∼2.0 × 10^6^ cells/mL. The cells were incubated for 72 h at 28 °C while shaking at 110 rpm. Cells were harvested by centrifugation at 1,000 × g for 20 min at 4 °C when cell viability was 80–85% and GFP fluorescence indicated >40% infection rate. The cell pellets were flash-frozen in liquid N_2_ and stored at −80 °C in lysis buffer containing 50 mM Bis-Tris HCl (pH 7.5), 300 mM NaCl, 0.5 mM TCEP, 0.05 mM EDTA, protease inhibitor cocktail tablets (cOmplete EDTA-free; Roche), and DNase I (Roche). For protein purification, cells were thawed and lysed by sonication (Branson Sonifier DS450). The lysate was clarified by ultracentrifugation at 50,000 × g for 60 min at 4 °C and filtered through 0.45 µm filters (Durapore Merck). The supernatant was loaded onto a Ni^2+^ affinity column (HisTrap HP 5 mL, Cytiva), washed with 50 mM Bis-Tris HCl (pH 7.5), 300 mM NaCl, 0.5 mM TCEP, 0.05 mM EDTA, 20 mM imidazole, and eluted with wash buffer supplemented with 350 mM imidazole (pH 8.0) directly onto a Strep-Tactin XT 4Flow 5 mL column (IBA). The final elution from the Strep-Tactin column was performed using BXT buffer (IBA). The eluted fractions were incubated with TEV protease for 16 h at 4 °C to remove the 10×His and Twin-StrepII affinity tags. The cleaved protein was separated from the tags by a second Ni^2+^ affinity chromatography step. Further purification was achieved by size-exclusion chromatrography on a HiLoad Superdex 200 16/600 pg column (Cytiva) equilibrated in 20 mM Hepes (pH 7.5) and 150 mM NaCl. The peak fractions were concentrated to 5–15 mg/mL and immediately used for crystallization experiments or flash-frozen in liquid N_2_ and stored at −80 °C for enzymatic assays.

The expression of the full-length BIN2, BIN2^K69R^ and BIN2^Y200F^ was done in *E. coli* BL21 (DE3) RIL cells grown in Terrific Broth containing 50 μg/mL kanamycin and 34 μg/mL chloramphenicol. When the culture reached an OD_600 nm_ of ∼0.8, protein expression was induced with 0.5 mM isopropyl β-D-1-thiogalactopyranoside (IPTG) at 18 °C for 16 h. Cells were harvested by centrifugation and resuspended in a buffer containing 50 mM Bis-Tris HCl (pH 7.5), 300 mM NaCl, 0.5 mM TCEP, and 0.1 mM EDTA, supplemented with a complete protease inhibitor cocktail tablet, DNase I, and 0.5 mg/mL lysozyme (Roth). Cell lysis was achieved by sonication. The lysate was then clarified by centrifugation at 50,000 × g for 30 min, and the supernatant was loaded onto a Ni ^2+^ affinity column (HisTrap HP 5 mL, Cytiva) equilibrated in 50 mM Bis-Tris HCl (pH 7.5), 300 mM NaCl, 0.5 mM TCEP, 0.1 mM EDTA, and 20 mM imidazole. After a washing step with the same buffer, BIN2 was eluted in buffer supplemented with 250 mM imidazole (pH 8.0). The 10×His-Streptavidin II-thioredoxin tag was cleaved overnight with TEV protease at 4 °C. The tag-free BIN2 protein was separated from the fusion tag by a second Ni^2+^ affinity chromatography step. Finally, BIN2 was applied to a HiLoad Superdex 200 16/600 pg column (Cytiva) equilibrated in 20 mM Hepes (pH 7.5) and 150 mM NaCl. The peak fractions containing BIN2 were concentrated, flash-frozen in liquid N_2_ and stored at −80 °C for enzymatic assays.

The human CDK1 – cyclin B1 – CKS1 complex, was co-expressed in Sf9 cells using baculoviruses as previously described (88). Typically, 25 ml of recombinant P3 baculoviruses were used to infect 500 ml of Sf9 insect cells at a cell density of roughly 2.0 × 10^6^ cells/mL. The cells were incubated for 48 h at 27 °C at 110 rpm, harvested at a cell viability rate of 80-85%, flash-frozen in liquid N_2_ and stored at −80 °C. The CDK1-cyclin B1-CKS1 complex was purified using StrepTactin Superflow Cartridge (Qiagen) columns. Lysis and wash buffer contained 50 mM Tris-HCl (pH 8.0), 500 mM NaCl, 10 mM β-mercaptoethanol and 5% (v/v) glycerol. The complex was eluted using wash buffer supplemented with 2.5 mM desthiobiotin and further loaded onto a HisTrap HP column (GE Healthcare Life Sciences). The complex was eluted from the HisTrap column in 50 mM Tris-HCl (pH 8.0), 500 mM NaCl, 10 mM β-mercaptoethanol, 5% (v/v) glycerol, and 250 mM imidazole. The CDK1-cyclin B1-CKS1 complex was loaded on a Superdex 200 Increase 10/300 GL column (Cytiva) size exclusion column as a final purification step. Peak fractions were pooled, concentrated and flash-frozen in liquid nitrogen and stored at −80 °C for enzymatic assays.

### Analytical size-exclusion chromatography

Gel filtration experiments were performed using a Superdex 200 Increase 10/300 GL column (Cytiva) pre-equilibrated in 20 mM Hepes pH 7.5, 150 mM NaCl. 80 μl of the respective protein was loaded sequentially onto the column, and elution at 0.75 ml/min was monitored by ultraviolet absorbance at 280 nm. Peak fractions were analyzed by SDS–PAGE gel electrophoresis.

### Protein crystallization and data collection

Crystals of the BSU1 Kelch domain developed in sitting drops composed of 0.25 μL of protein solution (BSU1^1-357^ at 15 mg/mL in 20 mM HEPES [pH 7.5], 150 mM NaCl) and 0.25 μL of crystallization buffer (18% [w/v] PEG 8,000, 5% [v/v] PEG 550 MME, 0.2 M sodium acetate trihydrate, 0.1 M sodium cacodylate trihydrate [pH 6.5]) suspended over 80 μL of the latter as reservoir solution. Crystals were transferred to reservoir solution supplemented with 20% (v/v) glycerol and snap-frozen in liquid N_2_. A complete native dataset to 2.25 Å was recorded at beam line X06DA of the Swiss Light Source, Villigen, Switzerland. Crystals of the BSU1 phosphatase domain grew in hanging drops composed of 1.5 μL of protein solution (BSU1^421-793^ at 5 mg/mL in 20 mM HEPES [pH 7.5], 150 mM NaCl) and 1.5 μL of crystallization buffer (0.7 M sodium citrate tribasic dihydrate, 0.1 M Tris [pH 8.0]) using microseeding protocols. Crystals were cryoprotected by serial transfer into 0.8 M sodium citrate tribasic dihydrate, 0.1 M Tris (pH 8.0), 20% (v/v) glycerol. A native dataset and a redundant single-wavelength anomalous dispersion (SAD) dataset close to the Zn K edge (9670.0 eV, three 360° wedges at 0.1° oscillation, with χ set to −15°, 0° and 15°) were collected to 2.1 and 2.7 Å resolution, respectively. Crystals of the BSU1 phosphatase-dead mutant (BSU1^421-793^ ^D510N^ ^H707K^ at 5 mg/mL in 20 mM HEPES [pH 7.5], 150 mM NaCl) developed in 1.6 M sodium / potassium phosphate (pH 7.0), were snap-frozen in mother liquor supplemented with 20% (v/v) glycerol, and diffracted to 1.6 Å resolution. Data processing and scaling was done with XDS (103).

### Structure solution and refinement

The structure of the BSU1 Kelch domain was solved by molecular replacement as implemented in in the program MoRDa (https://www.ccp4.ac.uk/morda-automatic-molecular-replacement-pipeline/). The highest scoring solution comprised one molecule of the Kelch domain of AtNSP1 (104) (PDB-ID pdb_00005gqt) in the asymmetric unit, and was used a starting model for automatic model building and refinement in ARP/wARP (105). The structure was completed in iterative rounds of manual model adjustment Coot (106) and restrained refinement in phenix.refine (107). The anomalous signal in the scaled SAD dataset from BSU1 phosphatase crystals extended to ∼4.5 Å resolution (108). 8 consistent Zn^2+^ sites were identified using XPREP/SHELXD (109). Site-refinement and phasing was done in autoSHARP (110), followed by non-crystallographic symmetry (NCS) averaging and density modification in phenix.resolve (111). The resulting starting phases were input in phenix.autobuild (112) and the structure was completed by manual model building in Coot (106) and restrained refinement in phenix.refine (107). The structure of phosphatase-dead BSU1 was solved by molecular replacement using the program Phaser (113) and using the refined BSU1 phosphatase structure as search model. Inspection of the final models with phenix.molprobity (114) revealed excellent stereochemistry (*SI Appendix*, Table S1). Structural representations were done with ChimeraX (115).

### *p*-NPP phosphatase assay

The phosphatase activity of BSU1^421-793^ (or of its catalytically dead variant BSU1^421-793^ ^D510N^ ^H707K^) toward *para*-nitrophenyl phosphate (*p*-NPP) was quantified by measuring the absorbance of the reaction product, *para*-nitrophenol (*p*-NP), at 405 nm using a plate reader (Tecan Infinite M Nano). Assays were performed in 100 µL reaction mixtures containing 50 mM Hepes (pH 7.5), 200 mM NaCl, and 2 mM MnCl₂ at 37 °C for 2, 4, or 8 min. At each time point, the reaction was quenched by mixing 50 µL of the reaction mixture with 20 µL of 150 mM EDTA. *p*-NPP substrate concentrations ranged from 0.5 to 20 mM, while the enzyme concentration was maintained at 5 µM. The quantity of released product was estimated using a standard curve of OD_405 nm_ versus *p*-NP standard concentrations. Enzyme-free blanks at OD_405 nm_ were measured at each time point to discard effect of non-enzymatic hydrolysis of *p*-NPP. The linearity of absorbance over time (R² > 0.95 for all conditions) confirmed that initial reaction rates were measured. The resulting kinetic data for BSU1^421-793^ were fitted to the Michaelis-Menten equation using RStudio (version 4.2.2) to determine the *V*_max_, *K*_M_, and *k*_cat_ values. All experiments were performed in triplicates (n = 3).

### Dephosphorylation assay of BIN2 at pTyr200

The protein phosphatase activity of BSU1^421-793^ toward phosphorylated Tyr200 (pTyr200) of BIN2 was assessed by immunoblotting. Assays were conducted in 100 µL reaction mixtures containing 50 mM Hepes (pH 7.5), 200 mM NaCl, and 2 mM MnCl₂ at 37 °C for 5, 10, 20, or 30 min. BIN2 and BIN2^Y200F^ were used at a concentration of 20 µM, resulting in a 4:1 substrate-to-enzyme ratio relative to either BSU1^421-793^ or to full-length BSU1^1-793^ (5 µM). Reactions were quenched at each time point by mixing 20 µL of the reaction mixture with 5 µL of SDS loading buffer (62.5 mM Tris-HCl pH 6.8, 30% glycerol, 0.01% [w/v] bromophenol blue), followed by incubation at 95 °C for 5 min. 2 µg of BIN2 and 0.5 µg of BSU1^421-793^ were resolved on a 10% BIS-Tris acrylamide SDS-PAGE gel using MES running buffer and transferred to a 0.2 µm PVDF membrane (IB34002, Thermo Scientific) using a semi-dry protocol (iBlot3, Thermo Scientific). Membranes were blocked with TBS-Tween (0.1%) containing 5% BSA for 1 hour at room temperature. For pTyr200 detection, membranes were probed for 1 h at room temperature with an anti-phospho-GSK3α/β (Tyr 279/216) monoclonal antibody (clone 5G-2F, mouse, Merck, ref: 05413, lot: 4229392) at a 1:2,000 dilution. Following three 10-min washes with TBS-Tween (0.1%), membranes were incubated for 1 h at room temperature with an anti-mouse-peroxidase polyclonal secondary antibody (goat, Promega, ref: W402B, lot: 0000169565) at a 1:3,000 dilution. Signal detection was performed using SuperSignal West Femto Maximum Sensitivity Substrate (34095, Thermo Scientific). The experiment was performed in triplicate (n = 3).

### Malachite green-based phosphatase assay

The protein phosphatase activity of BSU1^421-793^ (or its catalytically dead variant BSU1^421-793^ ^D510N^ ^H707K^) was quantified using the Malachite Green Phosphate Assay Kit (Sigma-Aldrich). The assay detects the release of inorganic phosphate produced by BSU1^421-793^ during the dephosphorylation of three synthetic 13-amino acid peptide substrates, RKLRRK(pT)GKRGSY, RKLRRK(pS)GKRGSY, and RKLRRK(pY)GKRGSY, which differ solely at the phospho-residue (designed based on Hein et al. (68); see *SI Appendix* Fig. S4). Synthetic peptides for the putative natural substrates BIN2 (193-KGEANIS(pY)ICSRFYR-207, BIN2^193-207^pY), CrCDKB1 (8-EKIGEG(pT)YGKVYK-20, CrCDKB^18-20^pT; or 8-EKIGEGT(pY)GKVYK-20, CrCDKB^18-20^pY) and BZR1 (168-<u>R</u>LRISN(pS)CPVTP<u>RY</u>-178, BZR1^168-178^pS) were based on previous reports (33, 34, 58, 24, 82, 83). All peptide were purchased from GenScript biotech. Reactions were performed in 100 µL mixtures containing 50 mM Hepes (pH 7.5), 200 mM NaCl, and 2 mM MnCl₂ at 37 °C. For RKLRRK(pT)GKRGSY, RKLRRK(pS)GKRGSY, and BZR1^168-178^pS, 50 nM BSU1^421-793^ were incubated with substrate concentrations ranging from 6.25 µM to 1000 µM for 2, 3, or 4 min. For RKLRRK(pY)GKRGSY, BIN2^193-207^pY, CrCDKB^18-20^pT and CrCDKB^18-20^pY, a higher BSU1^421-793^ enzyme concentration (2 µM) was used to ensure sufficient Pi release for malachite green reagent detection. Reactions were quenched by mixing 40 µL of the reaction mixture with 10 µL of malachite green reagent, and absorbance at 620 nm was measured after 25 min using a plate reader (Tecan Infinite M Nano). Enzyme-free blanks at OD_620 nm_ were measured at each time point to discard effect of non-enzymatic hydrolysis or background inorganic phosphate. The concentration of released inorganic phosphate was estimated using a standard curve of OD_620 nm_ versus inorganic phosphate standards. Kinetic parameters (*V*_max_, *K*_M_, and *k*_cat_) were calculated by fitting the data to the Michaelis-Menten equation in RStudio (version 4.2.2), with linear regression analysis confirming R² > 0.95 for all conditions. All experiments were conducted in triplicate (n = 3).

### Generation of transgenic Arabidopsis lines

For *bri1-5* suppression assay the BSU1 (At1g03445) or BSL2 (At1g08420) genomic fragments were amplified using Phusion High-Fidelity DNA Polymerase (New England Biolabs) according to manufacturer’s protocols and clone into a Golden Gate level 1 plasmid (116) (*SI Appendix*, Table S2). Point mutations were introduced using Gibson cloning (*SI Appendix*, Table S2). Level 2 binary plasmids were assembled providing the Cauliflower Mosaic Virus CaMV 35S promoter, the corresponding BSU1 version, a C-terminal 3xFlag tag, a NOS terminator and FastRed as selection marker. Final constructs were introduced into *Agrobacterium tumefaciens* strain GV3101 and then to the *bri1-5* mutant using the floral dip method (117). Transgenic T1 seeds were identified by mCherry fluorescence with a Zeiss Axio Zoom.V16 stereo microscope (mRFP filter) and a HXP200C illuminator. The *bsu1*, *bsu1 bsl1 bsl2* and *bsu1 bsl1 bsl3* mutants were generated by clustered regularly interspaced palindromic repeats (CRISPR/Cas9) gene editing (118). Guide-RNAs (gRNAs) were designed using the CRISPR-Pv2 website (http://crispr.hzau.edu.cn/CRISPR2/). Entry vectors containing the gRNAs under the control of the U6 promoter were generated by annealing and ligating forward and reverse primers. Then, gRNA entry vectors were used to create the binary vectors containing the *proU6*::*gRNA*, *proAtRPS5*::*Cas9* and FastRed selection marker using golden gate cloning. Constructs were introduced into *Agrobacterium tumefaciens* strain GV3101 and subsequently transformed into *A. thaliana* Col-0 ecotype using the floral dip method (117). Transgenic T1 seeds were identified via their fluorescent seed coats. Genotyping of CRIPR-Cas9 events was done by PCR followed by Sanger sequencing, Cas9 was segregated out in the T2 generation. Homozygous single *bsu1* and triple mutants *bsu1 bsl1 bsl2* and *bsu1 bsl1 bsl3* mutants were isolated in T4 generation by PCR and verified by Sanger-sequencing (Microsynth, AG). Quadruple segregating lines *bsu1 bsl1 bsl3 bsl2*+/− and *bsu1 bsl1 bsl2 bsl3*+/− were generated, but no quadruple homozygous lines could be obtained.

### Phenotypic analysis of BSU1/BSL2 over-expressing lines in *bri1-5*

20-25 transformants per line were selected, sterilized, put on half-strength Murashige and Skoog (^1/2^MS) medium containing 0.8% (w/v) agar plates and stratified at 4 °C for 48 h. Plates were then moved to a Percival growth chamber with 16 h / 8 h light / dark cycle at 22 °C for 7 d. After that, seedling were transferred to soil and kept in a controlled environment with 16h / 8h light / dark regime at 22 °C for 45 d.

### Western-blotting

To analyze the protein accumulation of BSU1-3xFlag and BSL2-3xFlag constructs, ∼100 mg of tissue sample were harvested and snap-frozen in liquid N_2_ in a 2 mL tube with glass beads. Samples were ground in a tissue lyzer (MM400, Retsch) and 300 µl of extraction buffer (50 mM Tris [pH 7.5], 150 mM NaCl, 1% [v/v] protease inhibitor cocktail [PE023, Merck]) was added to the tube. Samples were then gently agitated at 4 °C for 30 min in an orbital shaker (Intelli-Mixer RM-2M, ELMI). Next, samples were centrifugated at 10,000 × g for 20 min at 4 °C. 150 µL of supernatant was transferred to a new tube and 30 µl of 6x SDS sample buffer (0.5 M Bis-Tris [pH 6.6], 50% [v/v] glycerol, 6% [w/v] SDS, 0.5 M DTT with 0.05% [w/v] bromophenol blue) was added. Samples were boiled at 95 °C for 5 min and 30 μl were loaded and separated on a 10% SDS-PAGE gel. Proteins were transferred to a 0.2 μm PVDF membrane (IB34002, Thermo Scientific) using a semi-dry protocol (iBlot3, Thermo Scientific). Membranes were blocked for 1 h with 5% (w/v) milk powder in TBS-T (50 mM Tris [pH 7.5], 150 mM NaCl, 0.1% [v/v] Tween 20) at room temperature. Then membranes were incubated at room temperature for 2 h with Anti-FLAG M2-Peroxidase (HRP) antibody and a dilution of 1:5,000. Membranes were washed three times with TBS-T, detected using SuperSignal West Femto Maximum Sensitivity Substrate (Thermo Scientific) and visualized with Amersham Imager (Cytiva).

### Affinity purification coupled to mass spectrometry

The immunoprecipitation followed by mass-spectrometry (AP-MS) experiment was performed as previously described (119). Seedlings of *pro35S*::*BSU1^1-357^-3xFlag*, *pro35S*::*BSU1^421-793^-3xFlag* and *pro35S*::*BSU1^421-793^ ^D510N^ ^H707K^-3xFlag* were sterilized and plated on ^1/2^MS medium (Duchefa). Plates were stratified at 4 °C for 48 h and transferred to a Percival growth chamber with 16 h / 8 h light / dark cycle at 22 °C for 12 d. 100 mg of whole seedlings were harvested and snap-frozen in liquid N_2_. Samples were ground in a tissue lyzer (MM400, Retsch) and 300 µl of buffer was added (50mM Tris [pH 7.5], 150mM NaCl, 0.1% [v/v] Triton X-100, 10% [v/v] glycerol, 1% [v/v] protease inhibitor cocktail [PE023,Merck]). Tubes were gently agitated for 30 min at 4 °C and then centrifuged at 10,000 × g for 15 min at 4 °C. The supernatant was transferred to a new tube and centrifuged again at 10,000 × g for 15 min at 4 °C. Protein concentration was determined using Bradford method. All samples were adjusted to total protein concentration of 3 mg/mL. 200 µL of total protein was loaded to 50µl of buffer equilibrated Anti-FLAG magnetic beads (Sigma). Tubes were gently agitated for 30 min at 4 °C and then washed three times with 400 µL buffer. Proteins were eluted by adding 100 µL of 0.1 M glycine HCl (pH 3.0), with 5 min incubation. 10 µL were used for visualization on a silver-stained (Pierce™ Silver Stain for Mass Spectrometry, Thermo Scientific) SDS-PAGE gel. The remaining elution fractions were dried under speed-vacuum and proteins were digested with a mix of Trypsin/LysC using an iST kit (PreOmics). Peptides were then analyzed by nanoLC-ESI-MSMS using an easy-nLC 1000 liquid chromatography system (Thermo Fisher Scientific) coupled with a Q-Exactive HF Hybrid Quadrupole-Orbitrap Mass Spectrometer (Thermo Fisher Scientific). spectra were extracted and searched against the *Arabidopsis thaliana* reference proteome database (Uniprot, https://www.uniprot.org/, 2023-12, 27’448 entries) and an in-house database of common contaminant using Mascot (Matrix Science, London, UK; version 2.6.2). Trypsin was selected as the enzyme, with one potential missed cleavage. Precursor ion tolerance was set to 10 ppm and fragment ion tolerance to 0.02 Da. Carbamidomethylation of cysteine was specified as a fixed modification. Variable amino acid modification was oxidized methionine, as well as phosphorylation of serine, threonine and tyrosine. Peptide-spectrum matches were validated using Percolator a target FDR of 0.01 and a Delta Cn of 0.5. For label-free quantification, features and chromatographic peaks were detected using the “Minora Feature Detector” Node with the default parameters. PSM and peptides were filtered with a false discovery rate (FDR) of 1%, and then grouped to proteins with again a FDR of 1% (strict) or 5% (relaxed) and using peptides with high confidence levels. The ptmRS mode was used for phosphorylation modifications sites localization and validation. Only sites located with a probability of more than 75% were considered. Both unique and razor peptides were used for quantitation and protein abundances were calculated as the average of the three most abundant distinct peptide groups. The abundances were not normalized. “Protein abundance based” option was selected for protein ratio calculation and associated *p*-values were calculated with an ANOVA test (individual proteins).

### Yeast two-hybrid assays

BSU1^421-793^, BSU1^421-793^ ^M793R^ ^I750R^, BSL2 and MpBSLM coding sequences were synthesized (Twist Bioscience). MpCDK1, MpCKS, MpCYCB1, MpGCK and MpRBR were amplified from cDNA using high fidelity DNA polymerase KOD-plus (TOYOBO, No. KOD-201). All coding sequences were ligated to P6 and pB29 (Hybrigenics Services) providing an N-terminal GAL4 activating domain and a C-terminal LexA DNA binding domain, respectively, and the diploid TATA strain (Hybrigenics Services) was used (*SI Appendix*, Table S2). One colony of yeast was grown overnight at 30 °C with constant shaking at 200 rpm in 30 mL YPAD liquid media (20 g/L glucose, 20 g/L bacto-peptone, 10 g/L yeast extract and 0.04 g/L adenine hemisulfate). After 16 h, the culture was diluted to an OD_600 nm_ of 0.2 and grown until the OD_600 nm_ reached 0.4-0.6 before being transformed. Cells were collected at 1,000 × g for 5 min and re-suspended in 25 mL Tris-EDTA buffer (TE). Cells were centrifuged and re-suspended in 1xTE/LiAc to a final volume of 1.5 mL. 0.25 µg of prey plasmid, 0.25 µg of bait plasmid were mixed with 10 µl of herring sperm ssDNA (10 mg/ml). 100 µl of yeast cells and 600 ul of PEG/LiAc were added to each transformation, and vortexed for 10 s. The transformations were incubated 30 min at 30 °C while shaking at 200 rpm before heat shock at 42 °C for 15 min. After being cooled down on ice for 2 min the cells were spun down and re-suspended in 0.5 mL of TE. 100 µl of transformed cells were plated on SD plates (yeast nitrogen base with ammonium sulfate and without amino acids 6.7 g/L, glucose 20 g/L, bacto agar 20 g/L) without leucine and tryptophan and incubated at 30 °C for 3 d. Positive transformants were selected on SD – LW plates and growth overnight in selective SD liquid media in duplicates at 30 °C while shaking at 200 rpm. Each sample was diluted to an initial OD_600 nm_ of 0.2 and then sequentially diluted 10x (10x, 100x, 1,000x, 10,000x). 5 µl of each dilution was spotted on SD plates without leucine and tryptophan and with or without histidine. Plates were incubated at 30 °C for 2 d (Arabidopsis) or 3 d (Marchantia), respectively.

### Seedling hypocotyl growth assay

Seeds were surface sterilized, stratified at 4 °C for 2 d, and plated on ^1/2^MS medium containing 0.8% (w/v) agar and supplemented with 1 μM BRZ from a 10 mM stock solution in DMSO (Tokyo Chemical Industry) or with 0.1% (v/v) DMSO. To induce germination, seeds were exposed to light for 1 h and then incubated in the dark at 22 °C for 5 d. We then scanned the plates at 600 dots per inch resolution on a regular flatbed scanner (Perfection V600 photo; EPSON), measured hypocotyl lengths using Fiji (120) and analyzed the results in R version 4.1.2 (121) using the packages mratios (122) and multcomp (123). Rather than *p*-values, unadjusted 95% confidence limits for fold-changes are reported. A mixed-effects model for the ratio of a given line compared to the wild-type Col-0 is applied, allowing for heterogeneous variances to analyze log-transformed end point hypocotyl lengths. To evaluate treatment-by-mutant interactions, the 95% two-sided confidence intervals for the relative inhibition (Col-0: untreated versus BRZ-treated hypocotyl length)/(any genotype: untreated versus BRZ-treated hypocotyl length) are calculated for the log-transformed length.

### Seedling root growth inhibition assay

Seeds were sterilized, put on ^1/2^MS plates supplemented with 1% (w/v) sucrose and stratified at 4 °C for 48 h. Plates contained either 100 nM brassinolide (OlChemIm) from a 1 mM stock solution in 100% DMSO, or 0.01% (v/v) DMSO. Plates were transferred to a Percival growth chamber with 16 h / 8 h light / dark cycle at 22 °C for 7 d, after-which seedling were scanned (Perfection V600 photo; EPSON) and root length was manually measured in Fiji (120). Multiple comparisons of the genotypes vs. wild-type (Col-0) were performed using a Dunnett (124) test as implemented in the R package multcomp (123) (\**p* < 0.05; \*\**p* < 0.01; \*\*\**p* < 0.001).

### Stomatal index determination

Seeds were sterilized, put on ^1/2^MS plates supplemented with 1% (w/v) sucrose and stratified at 4 °C for 48 h. 7 d-old seedlings were used to determine stomatal indices, *scrm-D* (*125*) and *bsk1-2 bsk2-2* (126) were used as controls. For confocal imaging, seedlings were incubated in a 10 mg/L propidium iodide (PI) solution for 10 min and then washed five times with H_2_O. Cotyledon abaxial regions were imaged using a confocal LSM 780 optical microscope (Zeiss). PI staining was visualized with an excitation wavelength of 563 nm and emission was recorded between 571 nm and 656 nm, as previously reported (127). Mature stomata over a 0.5 mm by 0.5 mm epidermal surface area were counted manually using Fiji cell counter (120). A Dunnett test (124) was performed in the R package multcomp (123) to assess the statistical difference of the different genotypes compared to the Col-0 control (\**p* <0.05, \*\*\**p* < 0.01, \*\*\**p* < 0.001).

### Quantification silique and seed abortion phenotypes

Col-0, *bsu1 bsl1 bsl3*, *bsu1 bsl1 bsl2*, *bsu1 bsl1 bsl3 bsl2^+/−^* and *bsu1 bsl1 bsl2 bsl3^+/−^* plants were grown in soil for 5 weeks with a photoperiod of 16 h / 8 h at 22 °C. Representative pictures of inflorescence branch and siliques were taken (DS126761, Canon) and silique length were measured using Fiji (120) (n = 20). A Dunnett test (124) in the R package multcomp (123) was used to assess the statistical difference of the different genotypes compared to the Col-0 control (\**p* <0.05, \*\*\**p* < 0.01, \*\*\**p* < 0.001). Pictures from dissected siliques from the previously mention genotypes were taken with a binocular Nikon SMZ18 and seed abortion was evaluated on dissected siliques.

### β-glucuronidase (GUS) reporter assay

The β-glucuronidase (GUS) reporter activity analysis, 1154 bp upstream of BSU1 ORF were cloned to drive expression of *proBSU1*::*BSU1-GUS* and *proBSU1*::*GUS*, respectively (*SI Appendix*, Table S2). For BRI1 construct, 2073bp upstream of the BRI1 ORF (At4g39400) were used to generate the *proBRI1*::*GUS* reporter. All constructs were transformed into Ws-2 background. For each construct, 14-17 independent T2 lines were analyzed. Seeds were sterilized and stratified on ^1/2^MS plates for 48 h at 4 °C. Plates were then transferred to 22°C with 16 h / 8 h light dark cycle for 5 d, after-which staining was done for GUS activity in cotyledons. For 4^th^ leaf staining, seedlings were transferred to soil for additional 10 d before staining. Samples were incubated in ice-cold 90% (v/v) acetone for 20 min and rinsed with rinse solution (0.5 mM K_4_Fe(CN)_6_, 0.5 mM K_3_Fe(CN)_6_, and 50 mM NaPO_4_ buffer [pH 7.0]). Next, seedlings were vacuum infiltrated for 15 min in staining solution (0.5 mM K_4_Fe(CN)_6_, 0.5 mM K_3_Fe(CN)_6_, 10 mM EDTA, 0.1% (v/v) Triton X-100, 1 mM X-Gluc, and 100 mM NaPO_4_ buffer [pH 7.0]), and incubated at 37 °C for 12 h. Chlorophyll was removed by incubation in 70% (v/v) ethanol during several washing steps. Pictures were taken with a Nikon SMZ18 microscope.

### Callus induction

Seeds of the *proBSU1*::*GUS*, *proBSU1*::*BSU1-GUS* and *proBRI1*::*GUS* reporter lines were surface sterilized, stratified at 4 °C for 2 d, and plated on ^1/2^MS medium containing 0.8% (w/v) agar. Plates were transferred to a Percival growth chamber with 16 h / 8 h light / dark cycle at 22 °C. Seven-d-old seedlings were used as explant to induce callus formation. Cotyledon explants were transferred to B5 media with 2% (w/v) sucrose and supplemented with 2,4-D (500 μg/L) and kinetin (50 μg/L). Callus induction was done for 21 d at 22 °C in darkness, as previously reported (128). The GUS reporter assay was performed as described above.

### In vitro radioactive kinase assay

Phosphorylation of BSU1 by human CDK1 bound to cyclin B1 and CKS1 was conducted in 50 µL mixtures containing kinase buffer (50 mM Bis-Tris HCl, pH 7.5, 5 mM MgCl₂, 2 mM DTT), 10 µM ATP, 10 µCi of [γ-³²P]-ATP (6000 Ci/mmol, 10 mCi/mL) (Revvity), 10 nM of the CDK1 – cyclin B1 – CKS1 complex (88), and 10 µM BSU1^421-793^ substrate, or the BSU1^421-793^ ^T785A^ mutant protein, where Thr785 of the PxTPP motif is replaced by alanine. Reactions were incubated at 37 °C for 1, 5 and 10 min, respectively. Reactions were quenched by mixing 15 µL of the reaction mixture with 5 µL of SDS loading buffer (62.5 mM Tris-HCl, pH 6.8, 30% glycerol, 0.01% [w/v] bromophenol blue) followed by incubation at 95 °C for 5 min. Proteins were resolved on a 4–15% Mini-PROTEAN TGX Precast Gel (BioRad) using Tris-Glycine running buffer. The gel was then washed three times for 15 min each with a fixation solution (10% [v/v] acetic acid, 20% [v/v] ethanol), placed on a sheet of Whatman 3MM filter paper, covered with plastic wrap, and dried at 80 °C for 75 min under vacuum using a conventional gel dryer. The dried gel was exposed for 24 h to a phosphor screen for signal detection using the Sapphire FL Biomolecular Imager (Azure biosystems), with laser 638/424BPSP (n = 3).

The protein kinase activity of BIN2 toward BSU1^1-357^, BSU1^421-793^, and BSU1^1-793^ was assessed by observing the transfer of the radioisotope [γ-³²P] from ATP to BSU1 variants. Reactions were conducted in 50 µL mixtures containing kinase buffer (50 mM Bis-Tris HCl, pH 7.5, 5 mM MgCl₂, 2 mM DTT), 10 µM ATP, 10 µCi of [γ-³²P]-ATP (6000 Ci/mmol, 10 mCi/mL) (Revvity), 1 µM of the respective substrate BSU1 variant, and 30 nM BIN2. Reactions were incubated at 37 °C for 2 and 15 min (n = 3).

### Determination of in vitro phosphorylation sites

#### Sample preparation and protein digestion

Protein samples were loaded on a 12% mini polyacrylamide gel, stained by Coomassie Blue. Gel bands were excised and digested with sequencing-grade trypsin (Promega, Madison, WI, USA, Product N. V5111) or chymotrypsin (Promega, Prod.N. V1061), as described (129). Extracted tryptic peptides were dried and resuspended in 0.1% (v/v) formic acid, 2% (v/v) acetonitrile.

#### Liquid Chromatography-Mass Spectrometry analyses

LC-MS/MS analyses were carried out on a TIMS-TOF Pro (Bruker, Bremen, Germany) mass spectrometer interfaced through a nanospray ion source (“captive spray”) to an Ultimate 3000 RSLCnano HPLC system (Dionex). Peptides were separated on a reversed-phase custom packed 45 cm C18 column (75 μm ID, 100Å, Reprosil Pur 1.9 µm particles, Dr. Maisch, Germany) at a flow rate of 250 nl/min with a 2-27% acetonitrile gradient in 33 min followed by a ramp to 45% in 5 min and to 90% in 5 min (total method time: 65 min, all solvents contained 0.1% formic acid). Data-dependent acquisition was carried out using a standard TIMS PASEF method (130) with ion accumulation for 100 ms for each the survey MS1 scan and the TIMS-coupled MS2 scans. Duty cycle was kept at 100%. Up to 10 precursors were targeted per TIMS scan. Precursor isolation was done with a 2 or 3 m/z windows below or above m/z 800, respectively. The minimum threshold intensity for precursor selection was 2500. If the inclusion list allowed it, precursors were targeted more than one time to reach a minimum target total intensity of 20’000. Collision energy was ramped linearly based uniquely on the 1/k0 values from 20 (at 1/k0=0.6) to 59 eV (at 1/k0=1.6). Total duration of a scan cycle including one survey and 10 MS2 TIMS scans was 1.16 s. Precursors could be targeted again in subsequent cycles if their signal increased by a factor 4.0 or more. After selection in one cycle, precursors were excluded from further selection for 60s. Mass resolution in all MS measurements was approximately 35’000.

#### Data processing

MS data were analyzed using Mascot 2.8 (Matrix Science, London, UK) set up to search the *Arabidopsis thaliana* reference proteome based on the UniProt database, and a custom contaminant database containing the most usual environmental contaminants and enzymes used for digestion (keratins, trypsin, etc). Trypsin (cleavage C-terminal to K,R), allowing 2 missed cleavages and Chymotrypsin (cleavage C-terminal to Phe, Leu, Trp, Tyr, semispecific) were used as the enzyme definitions. Mascot was searched with a parent ion tolerance of 15 ppm and a fragment ion mass tolerance of 0.04 Da. Carbamidomethylation of cysteine was specified in Mascot as a fixed modification. Protein N-terminal acetylation, methionine oxidation and phosphorylation on Ser, Thr and Tyr were specified as variable modifications.

#### Data analysis

Scaffold (version Scaffold 5.0.0, Proteome Software Inc., Portland, OR) was used to validate MS/MS based peptide and protein identifications. Peptide identifications were accepted if they could be established at greater than 80.0% probability by the Percolator posterior error probability calculation (131). Protein identifications were accepted if they could be established at greater than 95.0% probability and contained at least 2 identified peptides. Protein probabilities were assigned by the Protein Prophet algorithm (132). All phosphopeptide identifications and phosphosite localizations reported were manually validated.

### Marchantia growth conditions

Wild type and CRISPR/Cas9-gene edited *Marchantia polymorpha* plants were Takaragaike-1 (Tak-1) males (133). Plants were asexually maintained and propagated through gemmae or by thallus stubs on ½Gamborg B5 medium (Sigma) adjusted to pH 5.5 with KOH, under constant LED-source white light (60 μmol/m^2^/s) at 22 °C on 90 mm square Petri dishes (Greiner) containing 0.8% (w/v) plant cell culture agar (Huber lab) as previously reported (134).

### Generation of stable transgenic *M. polymorpha* lines

Mp*bslm*^ge^ and Mp*gsk-3*^ge^ loss-of-function mutants were generated by CRISPR/Cas9 gene editing (118). gRNAs were designed using Casfinder (https://marchantia.info/tools/casfinder/). Primers were synthesized included forward 5’ CTCG– 3’ and reverse 5’ AAAC-3’ overhangs, respectively (*SI Appendix*, Table S2). Annealed primers were cloned into a pMpGE_En03 (Addgene #71535), plasmid digested with BsaI using T4 ligase (NEB). The entry gRNAs were subcloned into binary vector pMpGE010 (Addgene #71536) containing the Cas9 enzyme and a Hygromycin resistance gene as selection marker by Gateway Cloning, as described (135). The final plasmids were transformed into *Agrobacterium tumefaciens* strain GV2260 and plant transformations were done using the regenerated thallus protocol (133). Transgenic lines were genotyped by PCR using KOD polymerase (Merck) and by Sanger sequencing, using primers flanking the target regions. We selected lines bearing insertions or deletions that lead to frame shifts and early stop codons. The Mp*bslm*^ge^ Mp*gsk-4*^ge^ double mutant was generated by retransforming Mp*bslm*^ge^ with the gRNAs used to generate the Mp*gsk-3*^ge^ in plasmid pMpGE011 (Addgene #71537) providing a Chlorsulfuron resistance marker. The artificial micro-RNA was design in the amiRNA Design Helper tool (available at https://marchantia.info/tools/amir_helper/) and synthesized (Twist Bioscience). A fragment containing the miRNA was cloned by Golden-Gate into a vector providing a proMpEF1 promoter, a NOS terminator and hygromycin gene as selection marker. The final binary plasmid was transform into Tak-1 as described.

### Marchantia widefield microscopy

14 d-old Tak-1, amiRNA-MpBSLM, Mp*bslm*^ge^, Mp*gsk-3*^ge^ and Mp*bslm*^ge^ Mp*gsk-4*^ge^ plants were observed with an Axio Zoom V16 stereomicroscope (Zeiss) equipped with a 1x PlanNeoFluar NA 0.25 objective, an HXP 200 C fluorescence lamp (Kübler) and an Axiocam 512 colour CCD camera. Z stacks were acquired; bright-field images were acquired with episcopic white light, and fluorescence images were acquired with a DAPI filter (excitation 335-383 nm, emission 420-470 nm).

### Scanning electron microscopy (SEM)

14 d-old Tak-1, amiRNA-MpBSLM, Mp*bslm*^ge^, Mp*gsk-3*^ge^ and Mp*bslm*^ge^ Mp*gsk-4*^ge^ plants were transferred to aluminum holders on double-sided carbon tape (Electron Microscopy Sciences) and treated with a JFC-1200 gold sputter coater (JEOL). Imaging was performed with a JSM-6510LV scanning electron microscope (JEOL) in high vacuum mode with a spot size of 40 mm, an acceleration of 15 kV and a working distance of 12 mm.

### Marchantia cross section fixation, processing and histology

14 d-old Tak-1 and amiRNA-MpBSLM thalli and Mp*bslm*^ge^, Mp*gsk-3*^ge^ and Mp*bslm*^ge^ Mp*gsk-4*^ge^ thallus stubs were sectioned and fixed in phosphate buffer saline (PBS) pH 7.4 with 4% (v/v) formaldehyde and 0.2% (v/v) Triton X-100. The fixation was done overnight at 4 °C under agitation, after vacuum infiltration. Samples were then washed with PBS (2×15 min) and with H_2_O (2×10 min) before undergoing dehydration in a graded ethanol series (ethanol 30%, 50%, 70%, 90% and 100% [v/v] with 30 min incubation two times until the last bath in ethanol 100% [v/v] overnight at 4 °C). Technovit7100 (EMS, #14653) was prepared according to the manufacturer’s recommendations by supplementing it with hardener I, and samples were progressively infiltrated by incubation in 3:1, 1:1 and 1:3 mixes ethanol:Technovit7100 (each time 2 h under agitation at room temperature), before finally incubating in 100% Technovit7100 for 2 h at room temperature (after vacuum infiltration) and for another 40 h at 4 °C. Embedding was done in Technovit7100 supplemented with 1/15 Hardener II and 1/25 polyethylene glycol 400; polymerization was done for 30 minutes at room temperature followed by 30 min at 60 °C. Sectioning was performed with a Histocore AUTOCUT microtome (Leica) using disposable R35 blades and sections of 4 µm were deposited on SuperFrost slides. Sections were incubated 45 s in 0.2% (w/v) toluidine blue in dH _2_O, extensively rinsed with dH_2_O, then incubated one min in xylene and mounted in Pertex. The sections were observed with a Leica DM6B widefield microscope equipped with a DMC5400 CMOS camera (used with binning 2×2) and a 20x Fluotar NA 0.55 air objective.

### Image analysis

For a given dye, all images were acquired and post-processed identically. Image processing was done using Fiji (120). For toluidine blue pictures, tiling and stitching was done using the Leica LAS X Navigator tool. For wide-field microscopy images, a gamma of 0.45 was applied using Fiji, then focus stacking was performed with the Affinity Photo software (version 1.10.6.1665) and Fiji was finally used to apply a Cyan Hot look-up table to the fluorescence pictures and generate the overlay between brightfield and fluorescence.

### RNA-seq

Tak-1 and Mp*bslm*^ge^ plants were grown for 14 d in vitro on ½Gamborg B5 medium (Sigma) adjusted to pH 5.5 with KOH, under constant LED-source white light (60 μmol/m^2^/s) at 22 °C. For each biological replicate, 10 plants were pooled and RNA was extracted using the RNeasy plant mini kit (Qiagen). A Qubit fluorometer (Thermofisher) was used to determine RNA concentration, 100 ng per sample were analyzed to check quality control using 2100 Bioanalyzer system (Agilent Technologies). Library preparation and sequencing were performed by the iGE3 Genomic Platform at the Faculty of Medicine, University of Geneva (https://ige3.genomics.unige.ch/). Sequencing was performed with Novaseq 6000 machine from Illumina with 100 bp single-end output. Quality control of the reads and adaptor trimming were done with fastQC and Trimmomatic V6 (136). Genomic and transcript annotation files of the *Marchantia polymorpha* reference genome (137) and annotations were downloaded from MarpolBase (https://marchantia.info/download/MpTak_v6.1/). Read mapping to the reference was done with HISAT2 (138) (v2.2.1 with only the –dta option) and StringTie (139) (v2.2.1 with default options). DESeq2 (140) (v3.17) with default options as implemented in Galaxy server (https://usegalaxy.org/). Gene ontology enrichment analyses were performed in Panther (https://www.pantherdb.org/). DEGs annotation were manually curated to identify putative cell cycle regulated genes.

### EdU staining and imaging

EdU staining in Marchantia was performed using the Click-iT EdU (5-ethynyl-2′-deoxyuridine) Alexa Fluor 488 Imaging Kit (Thermo Fisher Scientific), as described (98). 3 d-old thalli of Tak-1 or Mp*bslm*^ge^ and Mp*gsk-3*^ge^ thallus stubs were grown in ^1/2^Gamborg’s B5 medium and soaked in half-strength liquid medium supplemented with 10 μM EdU at 22 °C for 3 h under continuous light. The samples were fixed in 50% (v/v) methanol / 10% (v/v) acetic acid (fixative solution) at 4 °C over night. After fixation, the samples were dehydrated through ethanol series of 80%, 90% and 100% (v/v) and a final 90% (v/v) acetone incubation at 80 °C for 5 min each, followed by fixative solution at 22 °C for 30 min. The samples were washed four times with 0.5% (v/v) Triton X-100 in PBS (137 mM NaCl, 2.7 mM KCl, 10 mM Na_2_HPO_4_, 1.8 mM KH_2_PO_4_), incubated with the EdU detection cocktail following the manufacturer’s instructions for 30 min in the darkness at room temperature, and rinsed four times with 0.5% (v/v) Triton X-100 in PBS. Finally, the samples were transferred to slides and incubated with chloral hydrate solution (25 g of chloral hydrate (C_2_H_3_Cl_3_O_2_) in 10 mL H_2_O) for 30 min. Images were obtained using a Leica DM6B widefield microscope equipped with a DFC9000GT sCMOS camera with excitation 450-490 nm and emission 500-550 nm.

### Confocal microscopy of Marchantia cell cycle reporter lines

Confocal laser scanning microscopy was done in 3 d-old Tak-1 thallus apical notches or in Mp*bslm*^ge^ and Mp*gsk-3*^ge^ thallus stubs transformed with the cell cycle reporter *proMpCYCD*::*Dbox-TdTomato* (96). Samples were transferred to slides with 10% (v/v) glycerol and image acquisition was performed on the SP8 microscope (Leica). TdTomato was excited at 552 nm, emission was recorded between 560 and 575 nm. Z-stack pictures were assembled using max intensity projection in Fiji (120).

## Data, Materials, and Software Availability

The atomic coordinates and structure factors have been deposited with the Protein Data Bank with accession codes pdb_00009TP5 (BSU1^1-357^ Kelch), pdb_00009TP7 (BSU1^421-793^ phosphatase domain) and pdb_00009TP8 (BSU1^421-793^ phosphatase-dead mutant). The mass spectrometry proteomics data have been deposited to the ProteomeXchange Consortium via the PRIDE partner repository with the dataset identifier PXD072713. Raw RNA-seq reads have been deposited with the NCBI sequence read archive (SRA; https://submit.ncbi.nlm.nih.gov/sra) with identifier <u>PRJNA1381677</u>.

## Acknowledgments

We thank Martin Bayer for providing *bsk1 bsk2* and *scrm-D* mutant seeds, and Andreas Boland, Silke Hauf, Martin Bayer, Roman Ulm and Philippe Rieu for their helpful comments on the manuscript and Zhiyong Wang for suggesting the BIN2 and CrBSL1 peptide substrates. We thank the University of Lausanne Protein Analysis Facility and the Proteomics Core Facility, Centre Medical Universitaire (CMU), University of Geneva for mass spectrometry analysis, and the staff of the Swiss Light Source (SLS) for technical assistance during data collection. This work was supported by the Swiss National Science Foundation grant 310030_205201 and by an International Research Scholar Award 55008733 by the Howard Hughes Medical Institute (to M.H.).

## Author Contributions

M.H. and A.M. designed the study. A.M. crystallized proteins, A.M. and M.H. solved and refined structures, P.R. and M.H. performed structural analyses, P.R. and A.M. performed phosphatase and kinase assays, J.Y. purified the CDK1 – cyclin B1 – CKS1 complex, O.P-T., L.B. and F.R. generated transgenic Arabiopsis lines, O.P-T., F.R. and H.C. performed Arabidopsis phenotyping experiments, O.P-T. and F.R. performed Y2H and GUS assays, O.P-T. performed the AP-MS experiment, F.R. generated transgenic Marchantia lines, performed phenotyping experiments, RNA-seq, and together with H.C., C.F. and S.L. performed and analyzed the SEM, histology and microscopy experiments. L.A.H. performed statistical analyses. M.H. was responsible for conceptualization, supervision, funding acquisition, and project administration. M.H., P.R. and F.R. wrote the original draft. All authors reviewed and edited the manuscript.

## Competing Interest Statement

The authors declare that they have no competing interests.

## Classification

Biological sciences, Plant Biology and Biochemistry

## Supporting Information for

**Fig. S1.**
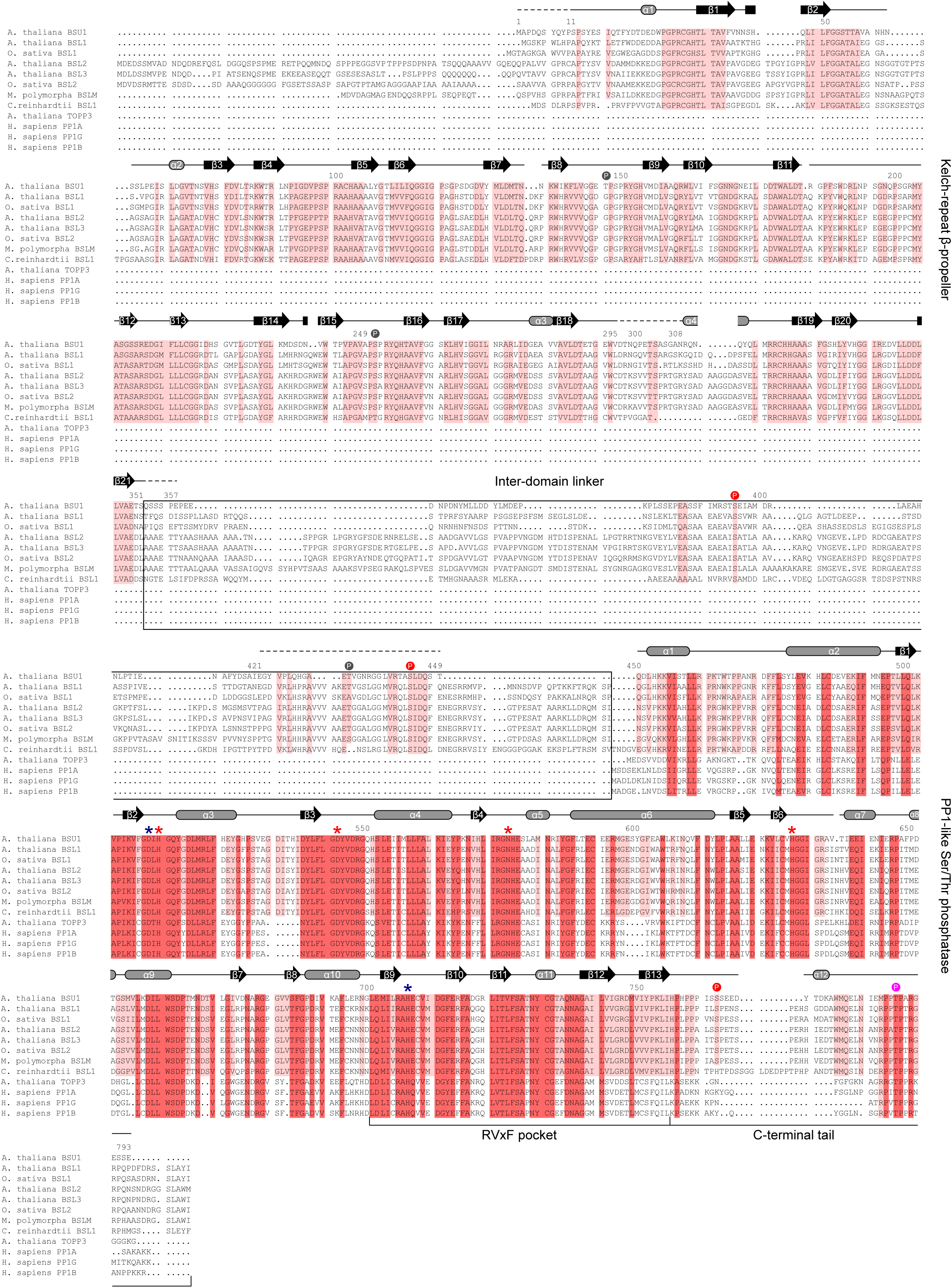
Structure-based sequence alignment of BSU1-related protein phosphatases. Shown is a multiple sequence alignment (*A. thaliana* BSU1, https://uniprot.org UniProt-ID: Q9LR78; *A. thaliana* BSL1, Q8L7U5; *O. sativa* BSL1, Q60EX6; *A. thaliana* BSL2, Q9SJF0; *A. thaliana* BSL3, Q9SHS7-1; *O. sativa* BSL2, Q2QM47; *M. polymorpha* BSLM, A0A2R6VWR4; *C. reinhardtii*, phytozome.org (v.5.6) identifier Cre01.g050850; *A. thaliana* PP1/TOPP3, P48483; *H. sapiens* PP1α, P62136; *H. sapiens* PP1γ, P36873; *H. sapiens* PP1β, P62140) and including a BSU1 secondary structure assignment (α-helices in gray, β-strands in black) calculated with the program DSSP (143). Conserved residues among plant Kelch-phosphatases are highlighted in light red, residues also conserved among PP1-type Ser/Thr phosphatases are shown in dark red, residues involved in metal ion coordination are denoted by a red *, the position of the D510N and H707K mutations are highlighted in blue. Phosphorylation sites are highlighted by a colored circle in either red (CDG1-mediated phosphorylation), magenta (CDK1-mediated phosphorylation) or gray (BIN2-mediated phosphorylation).

**Fig. S2.**
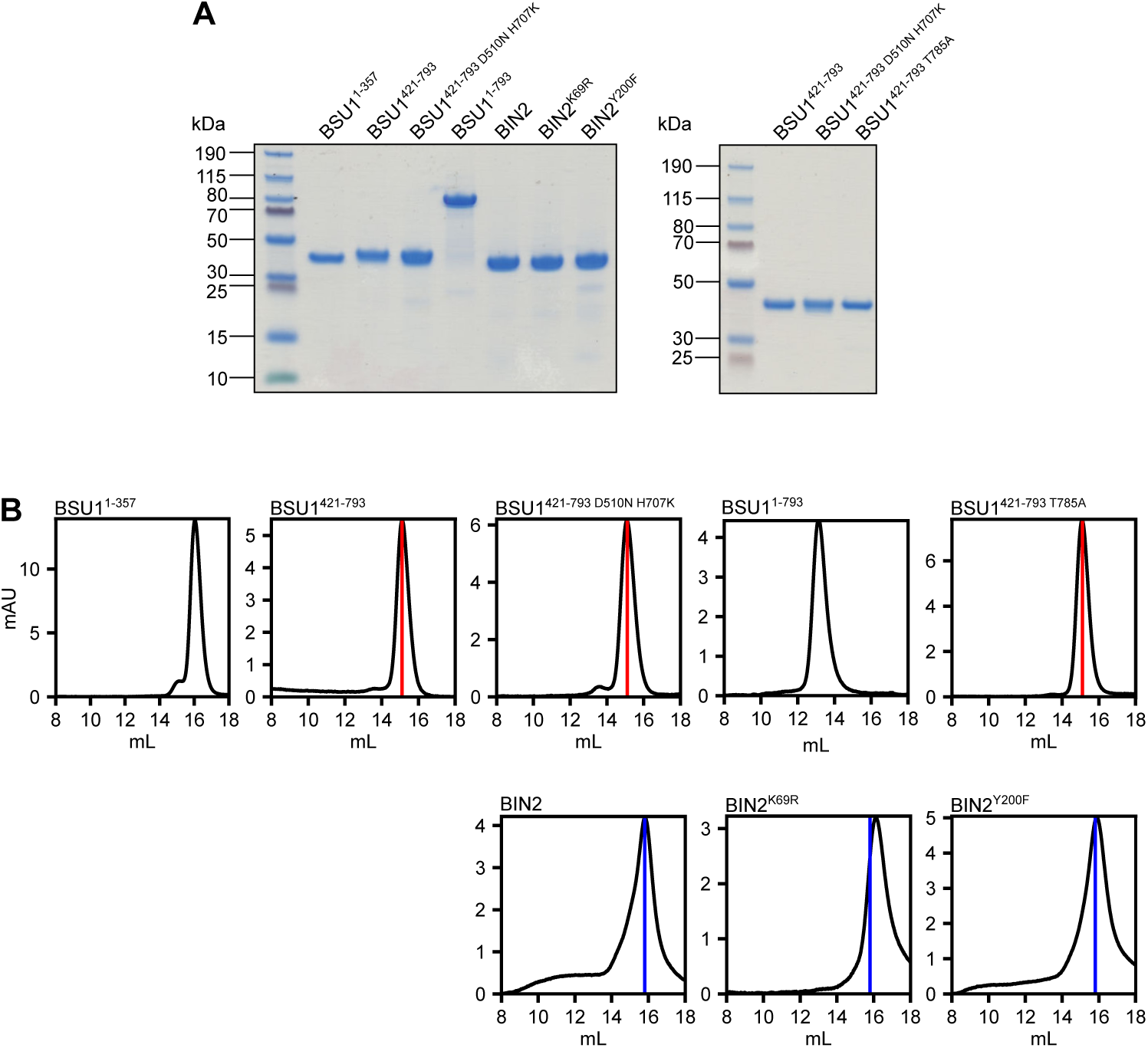
Purity of the BSU1 and BIN2 preparations used in this study. (*A*) Coomassie-stained SDS-PAGE analysis of the different wild-type and mutant fragments of BSU1 and BIN2. (*B*) Analytical size exclusion chromatography analyses of wild-type and mutant BSU1 and BIN2 fragments. 80 µL of sample were loaded to a Superdex 200 increase HR10/30 column and sample elution at 0.75 mL/min was monitored by UV absorption at 280 nm. The red and blue vertical lines indicate the characteristic elution volumes for monomeric wild-type BSU1^421-793^ and full-length BIN2 proteins, respectively.

**Fig. S3.**
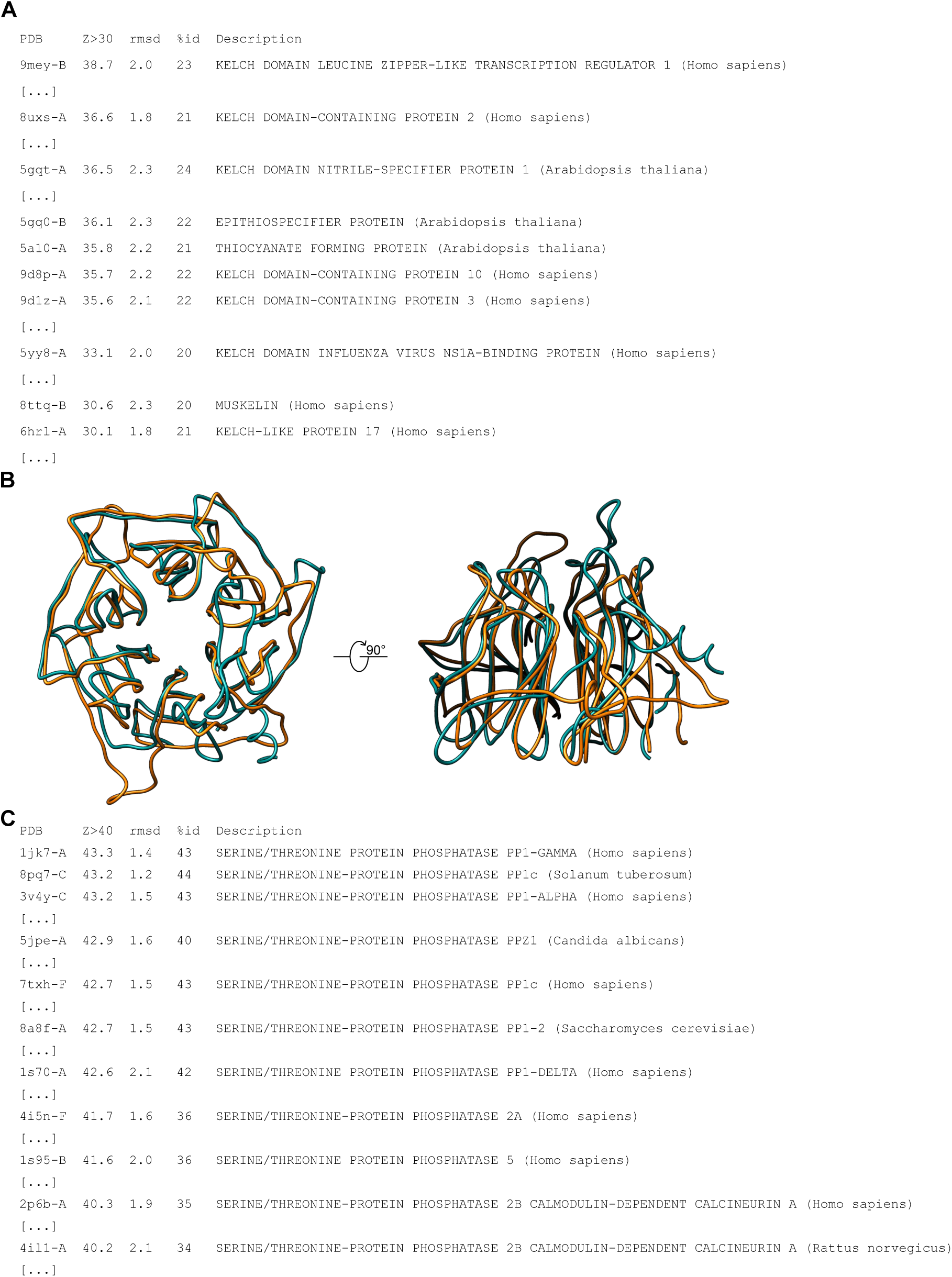
BSU1 shares structural homology with Kelch domain-containing proteins and with PP1-like Ser/Thr phosphatases. (*A*) Selected top hits from the DALI webserver (http://ekhidna2.biocenter.helsinki.fi/dali/) (64) using the isolated BSU1^11-351^ Kelch domain as search model. (*B*) Structural superposition of the BSU1 Kelch domain (cyan C_α_ trace) with the Kelch domain of human Leucine Zipper-like Transcription Regulator 1 (in orange, PDB-ID: pdb_00009mey, r.m.s.d. ∼1 Å comparing 184 corresponding C_α_ atoms) (60). (*C*) Selected DALI hits for the BSU1^450-793^ phosphatase domain.

**Fig. S4.**
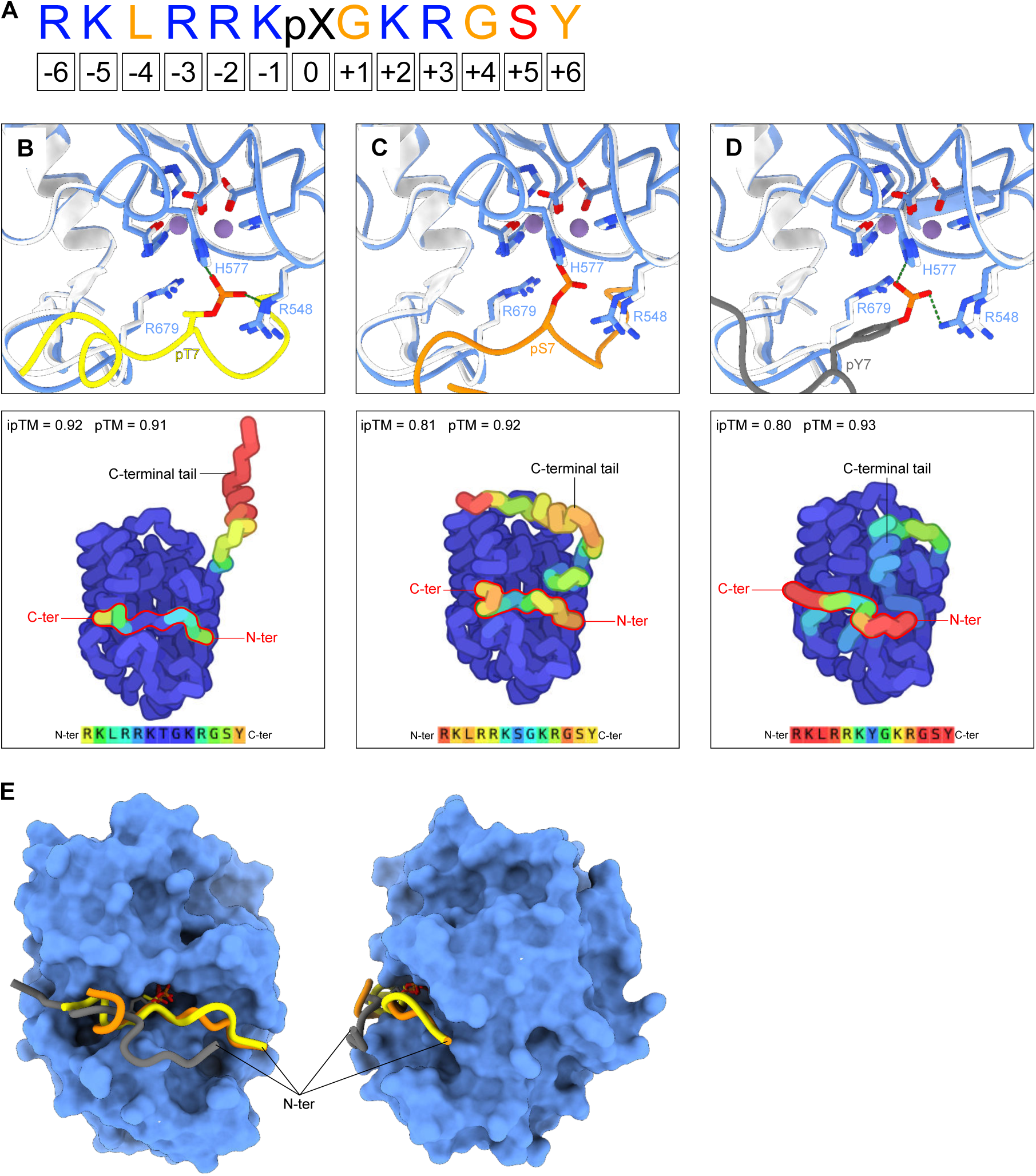
Design and molecular docking of BSU1 synthetic substrate peptides. (*A*) Amino acid sequence of the synthetic peptide used in BSU1 phosphatase substrate specificity assays. Basic, acidic, and hydrophobic residues are highlighted in blue, red, and yellow, respectively. The central phosphorylated residue (pThr, pSer, or pTyr) is denoted as pX. Residue positions relative to pX are indicated below, with negative and positive numbers representing upstream and downstream residues, respectively. Based on the analysis by Hein et al. (68), PP1 phosphatases exhibit a preference for basic residues at positions −6, −5, −3, −2, −1, +2, and +3, while there is no amino acid preference at position +1, except for the exclusion of proline when pSer is present. Positions +4, +5, and +6 show no preference; thus, a Gly-Ser-Tyr linker was incorporated at the C-terminus to accurately estimate the molar concentration of the peptide in solution, using the molar extinction coefficients at A_280 nm_. (*B*-*D*) AlphaFold3-predicted complex structures of BSU1^421-793^ bound to Mn^2+^ ions and three distinct phosphopeptides: the pThr-containing peptide in *B*, the pSer-peptide in *C*, and the pTyr-peptide in *D*, as implemented in https://alphafoldserver.com/ (69). The upper panels provide a close-up view of BSU1 active site in ribbon representation. The experimental crystal structure of BSU1^421-793^ is shown in white, the AlphaFold3 docking models are in blue. Phosphopeptides are colored in yellow (pThr), in orange (pSer) and in dark gray (pTyr). Predicted salt bridges between the phosphate group and BSU1 residues are depicted as dotted lines (in green). The lower panels present an overview of the AlphaFold3 docking models as C_α_ chain traces, colored by predicted local distance difference test (pLDDT) score (very high, dark blue, pLDDT > 90; high, cyan, 90 > pLDDT > 70; low, yellow, 70 > pLDDT > 50; very low, red, pLDDT < 50) and visualized using Py2DMol (https://py2dmol.solab.org). AlphaFold3 pTM and ipTM values are reported alongside (69). (*E*) Molecular surface representation of BSU1^421-793^ (in blue) bound to the different phosphopeptides, colored as in panels *B*-*D*. Note that the pTyr-containing peptide can only be accommodated in the active site adopting a highly unusual twisted conformation.

**Fig. S5.**
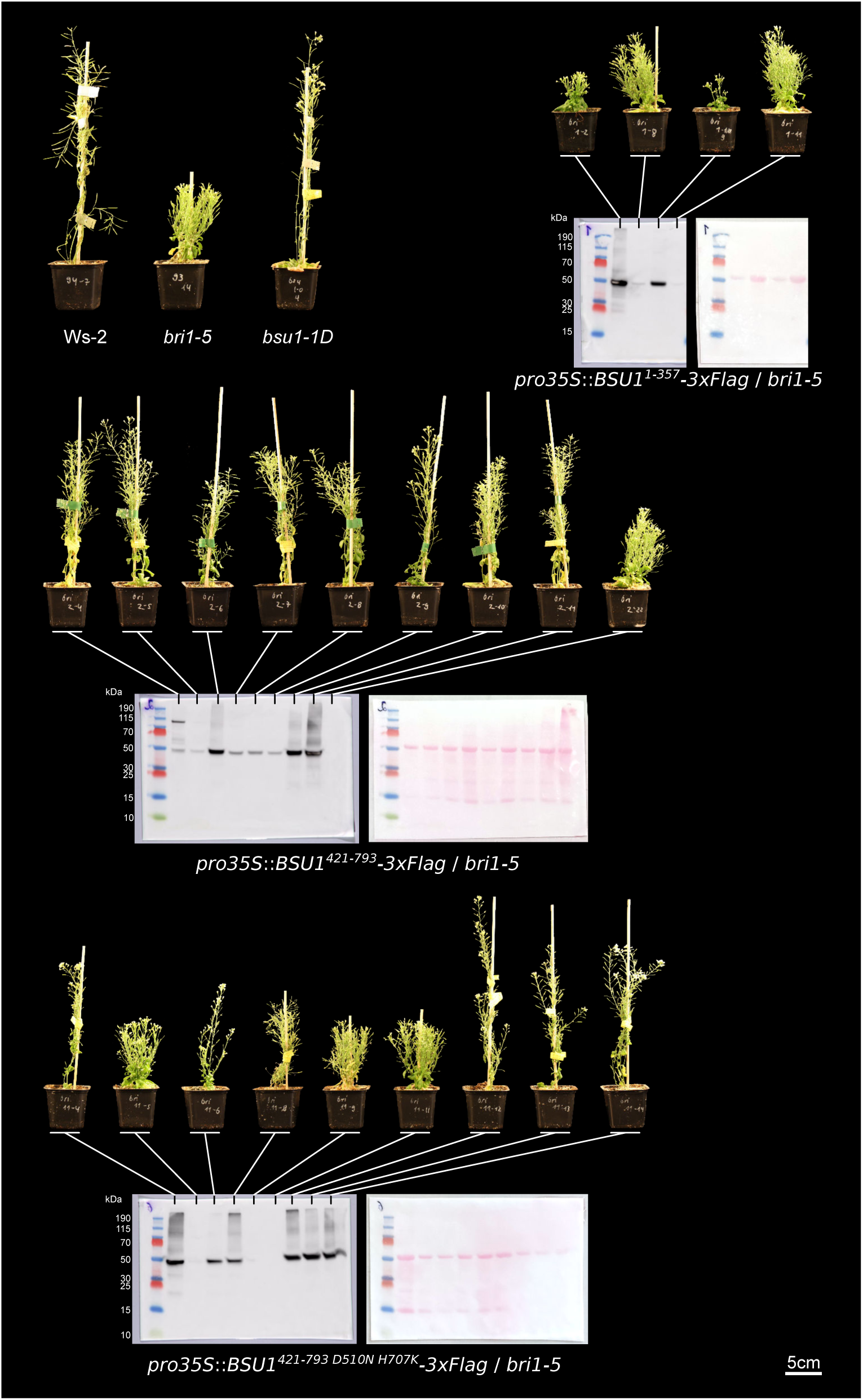
The isolated BSU1 Kelch domain further enhances the growth phenotype of the *bri1-5* mutant. Shown are shoot growth phenotypes of 45 d-old *bri1-5* plants expressing BSU1^1-357^ (Kelch domain), BSU1^421-793^ (phosphatase domain) or BSU1^421-793^ ^D510N^ ^H707K^ (phosphatase-dead) variants from the 35S promoter. Shown are T1 generation plants, the Ws-2 wild type, *bri1-5* (142) and the *bsu1-1D* (36) allele. BSU1 protein expression was assayed by western blot using an α-Flag antibody, the Ponceau-stained membranes are shown on the left as loading controls.

**Fig. S6.**
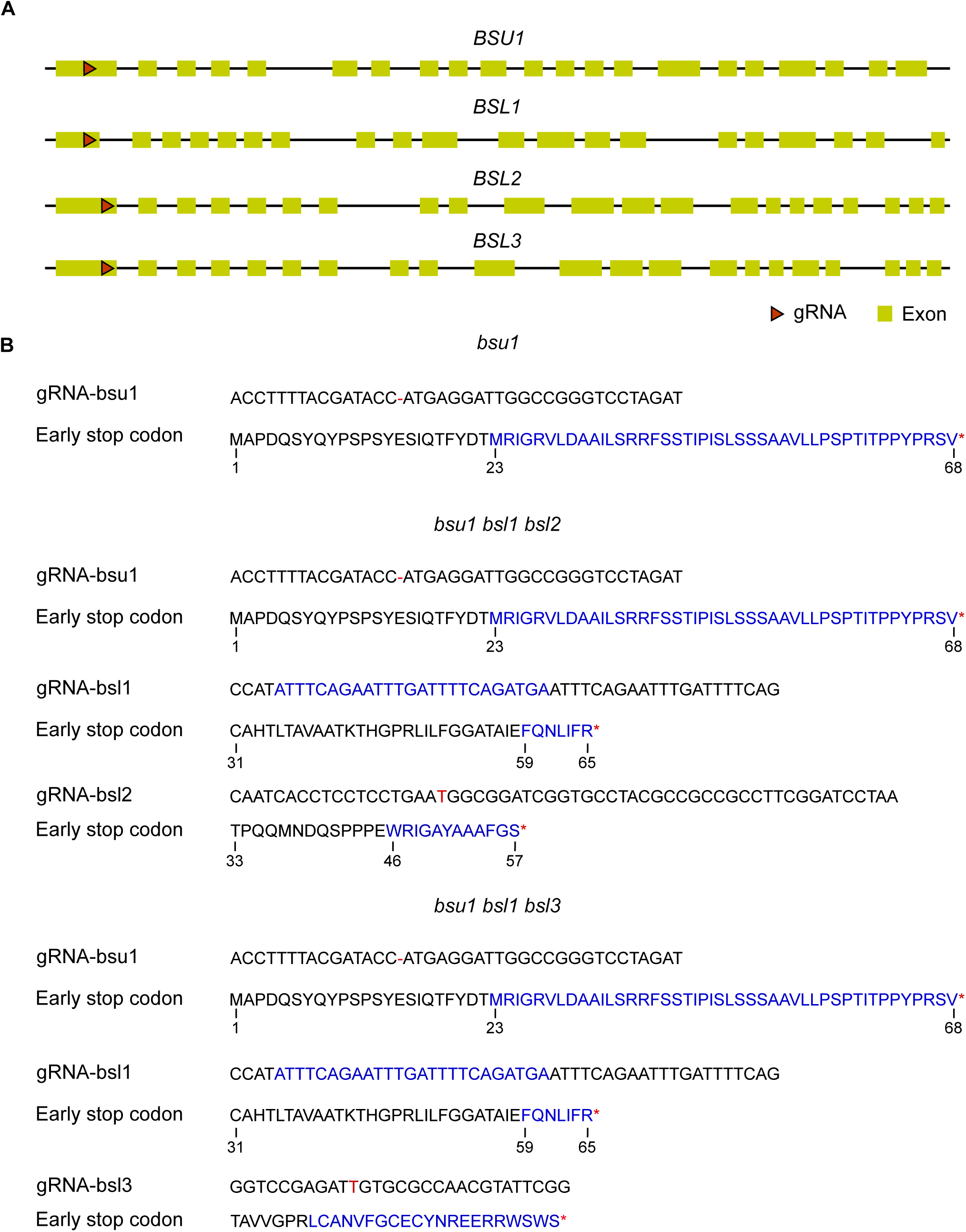
Arabidopsis *bsu1* and *bsl1-3* mutant CRISPR/Cas9 gene editing events. (*A*) Genomic representations of BSU1 (TAIR-ID: At1g03445), BSL1 (At4g03080), BSL2 (At1g08420) and BSL3 (At2g27210) genes with exons in yellow and introns shown as black lines. sgRNA targeting regions are represented with red arrowheads. (*B*) CRISPR events are shown below the corresponding genotype, as confirmed by Sanger sequencing. The altered DNA sequence and the stop codon are shown in red, the corresponding amino acid sequence is shown in black (wild type) and blue (altered sequence by the frame shift editing event).

**Fig. S7.**
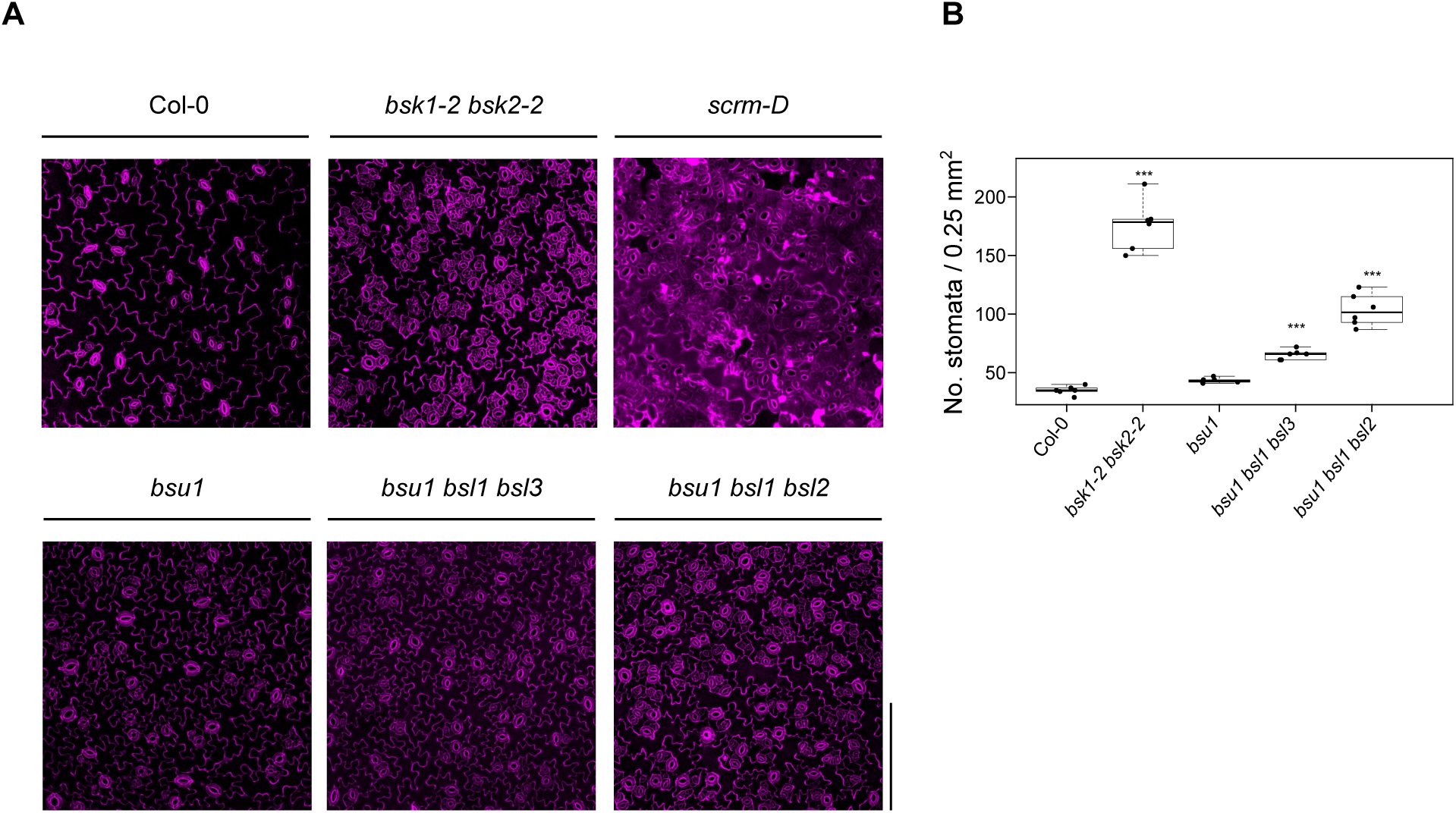
Deletion of multiple Kelch phosphatase triggers stomata over-proliferation. (*A*) Representative confocal images of 7 d-old abaxial cotyledon regions for Col-0, *bsk1-2 bsk2-2* (126), *scrm-D* (125), *bsu1* and different *bsu1 bsl* mutant combinations, stained with PI (in magenta). (scale bar = 200 µm). (*B*) Stomatal index quantification represented in box plots (n = 6; bold black line, median; box, inter-quartile range (IQR); whiskers, lowest/highest data point within 1.5 IQR of the lower/upper quartile, raw data shown as individual points) and including Dunnett-type (124) two-sided multiplicity-adjusted *p*-values for the comparison against the Col-0 control (\**p* < 0.05; \*\**p* < 0.01; \*\*\**p* < 0.001). The stomatal index of the strong *scrm-D* allele could not be determined reliably.

**Fig. S8.**
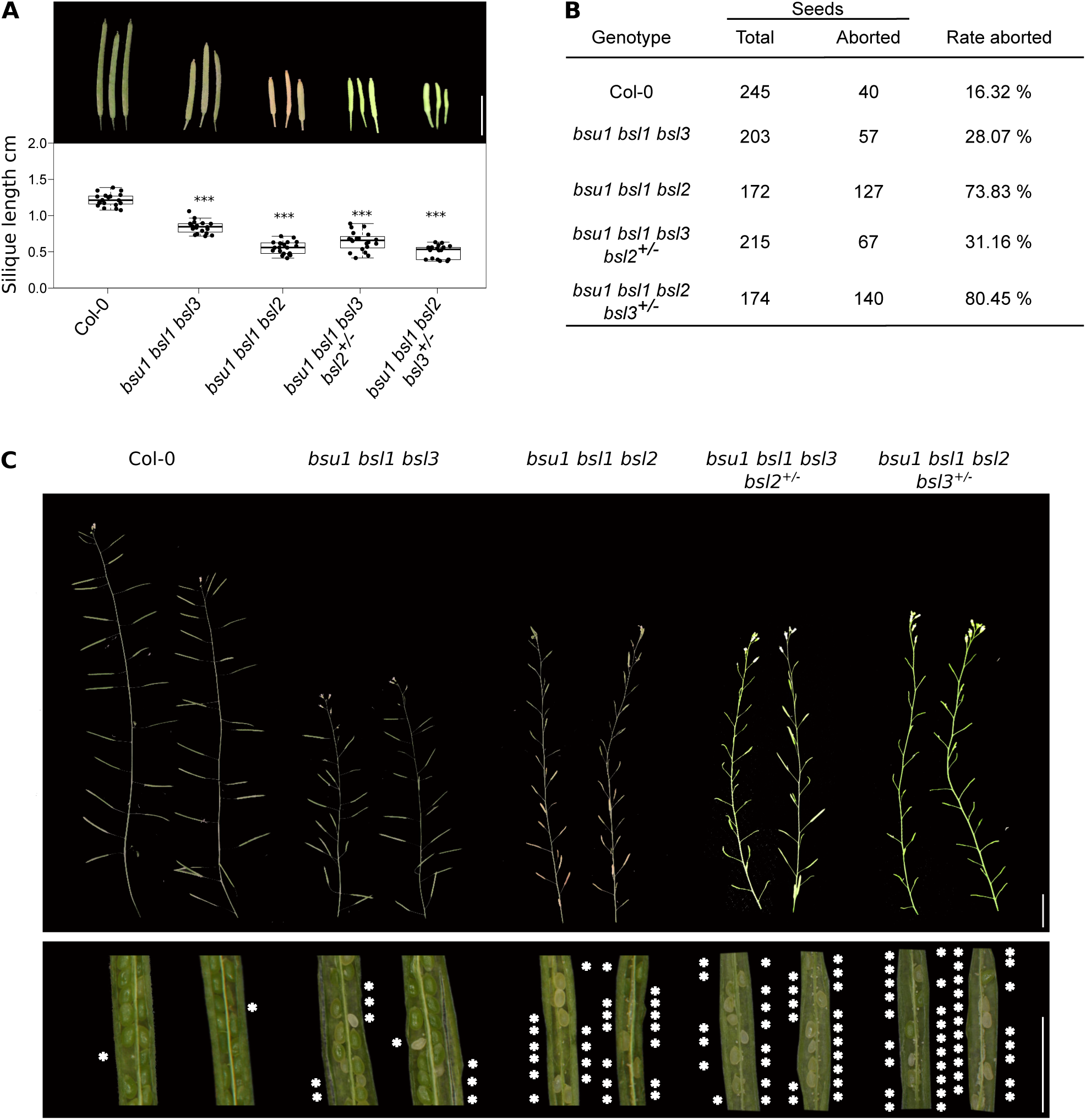
Higher order *bsu1 bsl* mutants have fertility defects. (*A*) Quantification of silique length in different *bsu1 bsl* mutant combinations with representative siliques shown on top (scale bar = 0.5 cm) and quantifications below (n = 20). Box plot bold black line, median; box, inter-quartile range (IQR); whiskers, lowest/highest data point within 1.5 IQR of the lower/upper quartile, raw data shown as individual points. Dunnet-type (124), two-sided multiplicity-adjusted *p*-values for the comparison against the Col-0 control are shown alongside (\**p* < 0.05; \*\**p* < 0.01; \*\*\**p* < 0.001). (*B*) Quantification of seed abortion in Col-0 and different *bsu1 bsl* mutant combinations. The table summarizes the total number of seed and aborted seeds. The abortion rate is expressed as percentage of wild type seeds. Shown below in (*C*) are representative shoot branches of the different genotypes on the top (scale bar = 1 cm) and individual opened siliques below (scale bar = 0.2 cm). Aborted seeds are denoted by white asterisks.

**Fig. S9.**
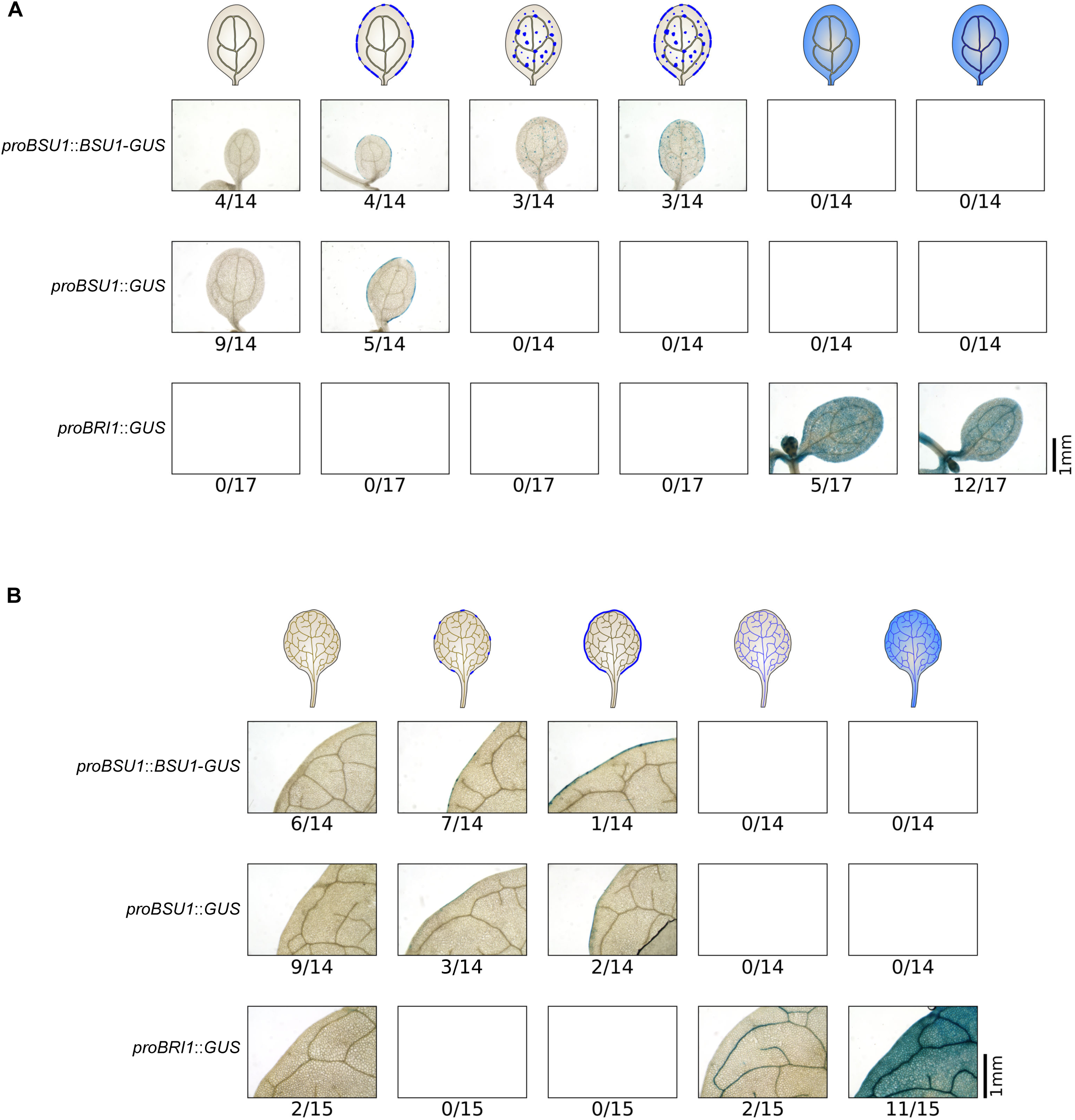
The expression patterns of BSU1 and BRI1 are not overlapping. Shown are spatiotemporal expression patterns of BSU1 and BRI1 reporter lines determined by β-glucuronidase activity (GUS) assays. A sequence ∼1kb upstream of the BSU1 ORF was used to drive *proBSU1*::*BSU1-GUS* and *proBSU1*::*GUS*, respectively. A ∼2kb upstream of BRI1 ORF was used to drive *proBRI1*::*GUS*. 14-17 independent T2 lines for each construct were stained for GUS activity, either using 5 d-old cotyledons (*A*), or the 4th leaf of 15 d-old soil-grown plants (*B*). Examples for the different categories summarized in the star plots in Fig. 2*F* are represented.

**Fig. S10.**
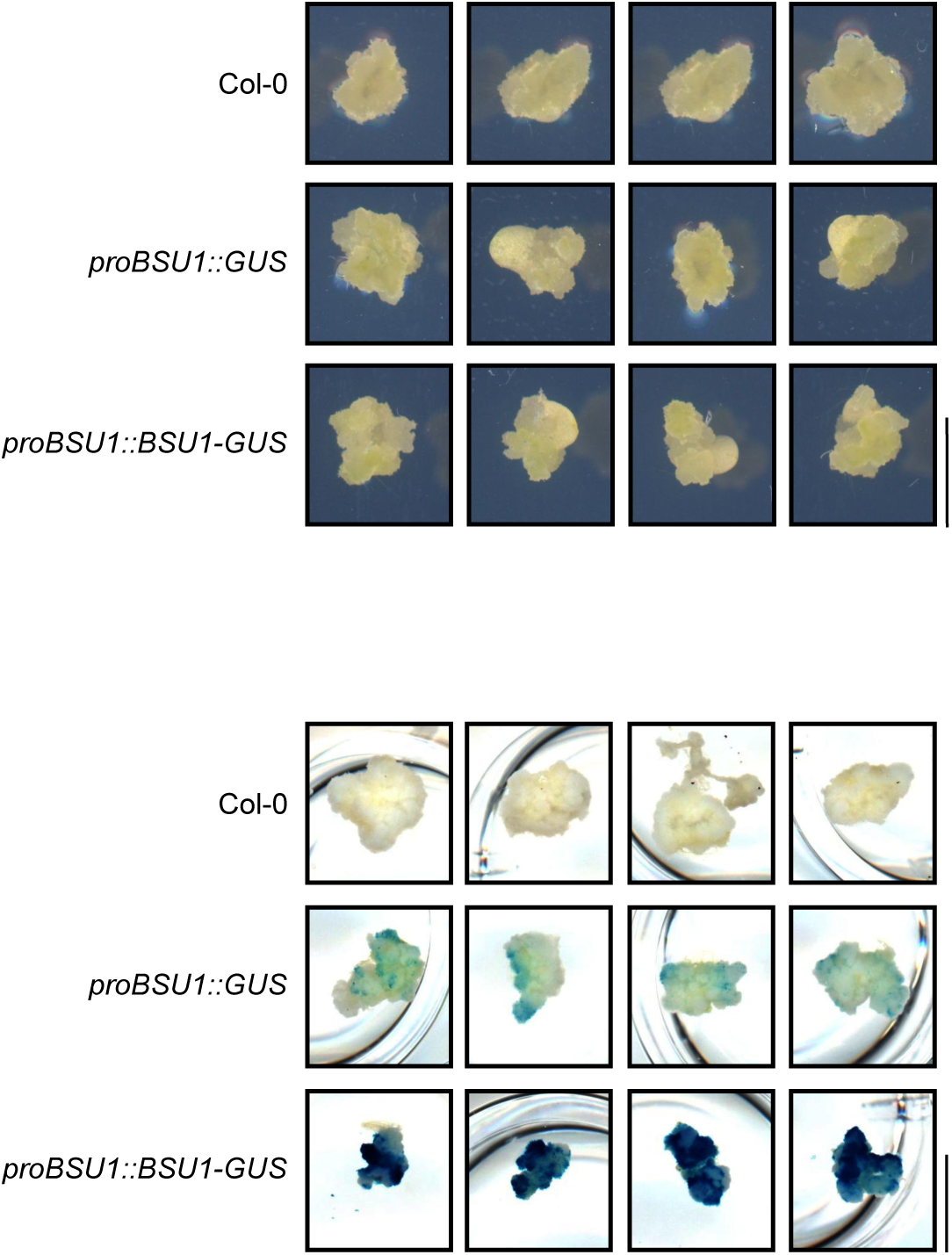
BSU1 expression pattern during callus induction. Promoter β-glucuronidase (GUS) reporter assay for 3-week-old callus induced from Col-0, *proBSU1*::*GUS* and *proBSU1*::*BSU1-GUS* plants. Representative pictures of the induced calli (top panel) and stained calli (bottom panel) are shown for all genotypes (scale bar = 0.5 cm).

**Fig. S11.**
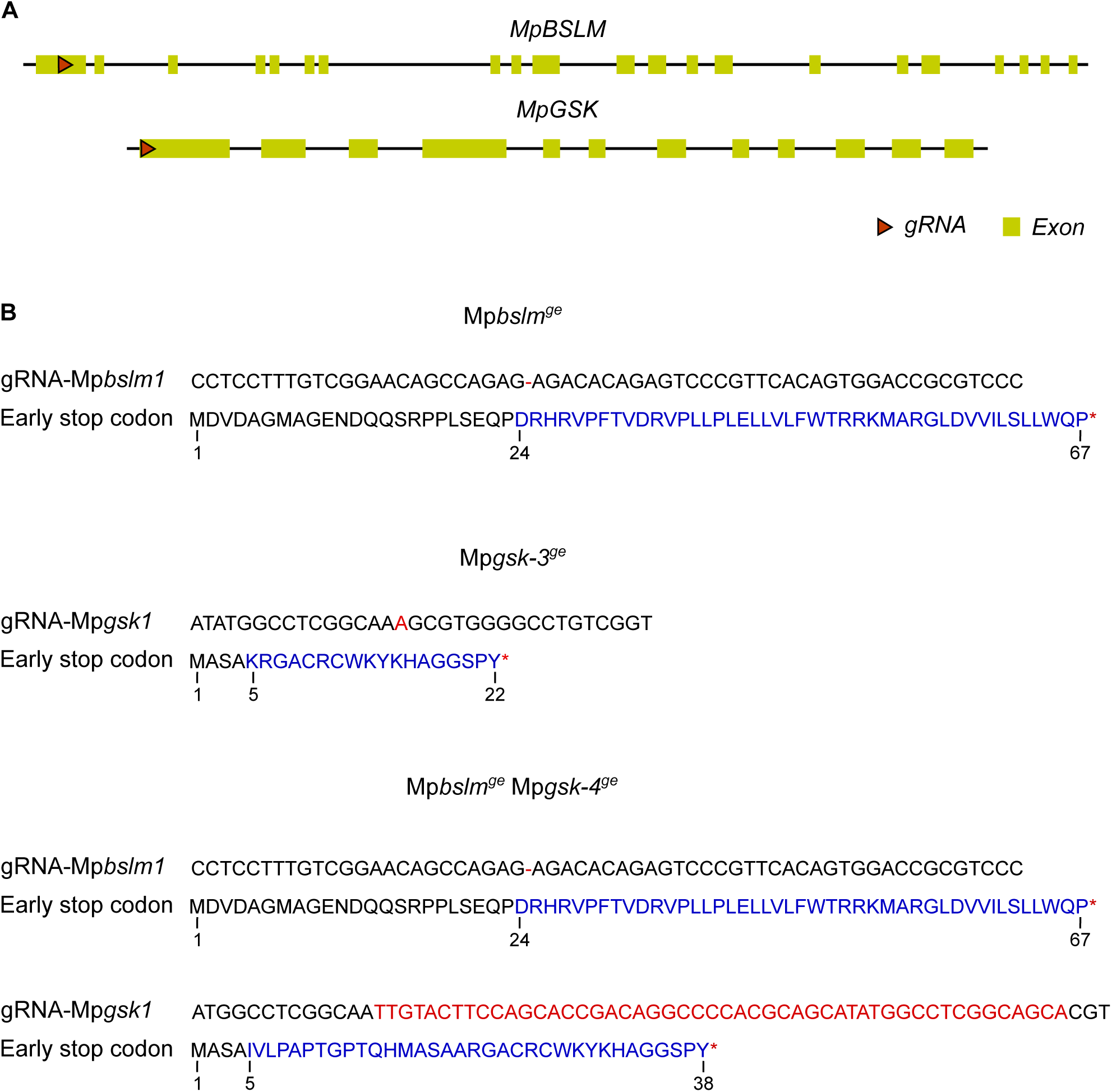
Marchantia Mp*bslm*^ge^ and Mp*gsk*^ge^ mutant CRISPR/Cas9 gene editing events. (*A*) Genomic representations of MpBSLM (MpVg00440, https://marchantia.info) and MpGSK (Mp7g04170) genes with exons in yellow and introns shown as black lines. SgRNA targeting regions are represented with red arrowheads. (*B*) CRISPR events are shown below the corresponding genotype, as confirmed by Sanger sequencing. The altered DNA sequence and the stop codon are shown in red, the corresponding amino acid sequence is shown in black (wild type) and blue (altered sequence by the frame-shift editing event).

**Fig. S12.**
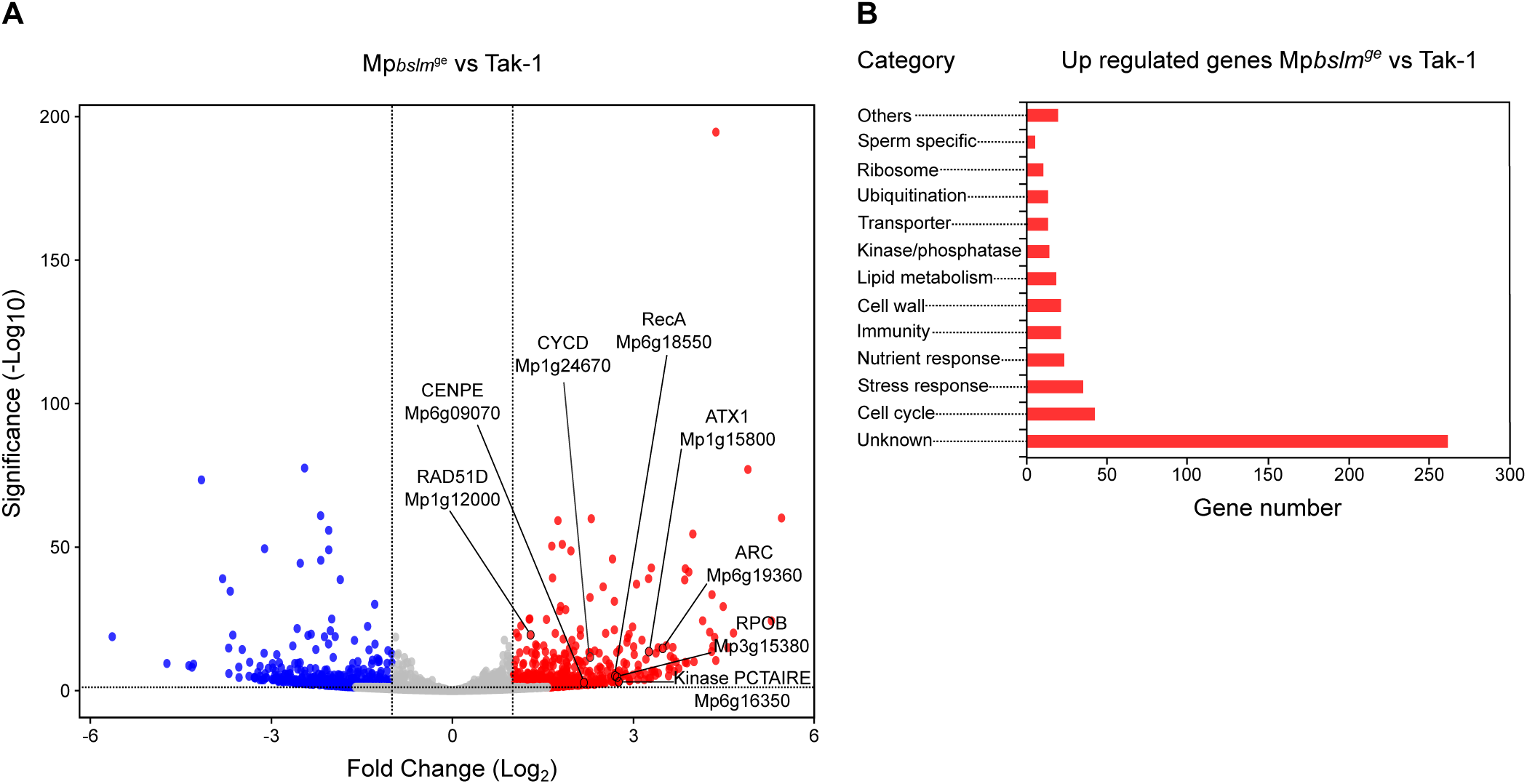
Summary of a RNA-seq experiment comparing 14 d-old Mp*bslm*^ge^ mutant plants with the Tak-1 wild type. (*A*) Volcano plots showing differentially expressed genes (DEGs) (Log_2_FC > 1, in red and Log_2_FC < –1, in blue) of Mp*bslm*^ge^ vs. Tak-1. Putative cell cycle-regulated genes discussed in the text are indicated in black (*B*) Manually curated gene-ontology classification of DEGs. Up-regulated genes with |log_2_(FC)|>1 and *p* < 0.05 were considered differentially expressed.

**Fig. S13.**
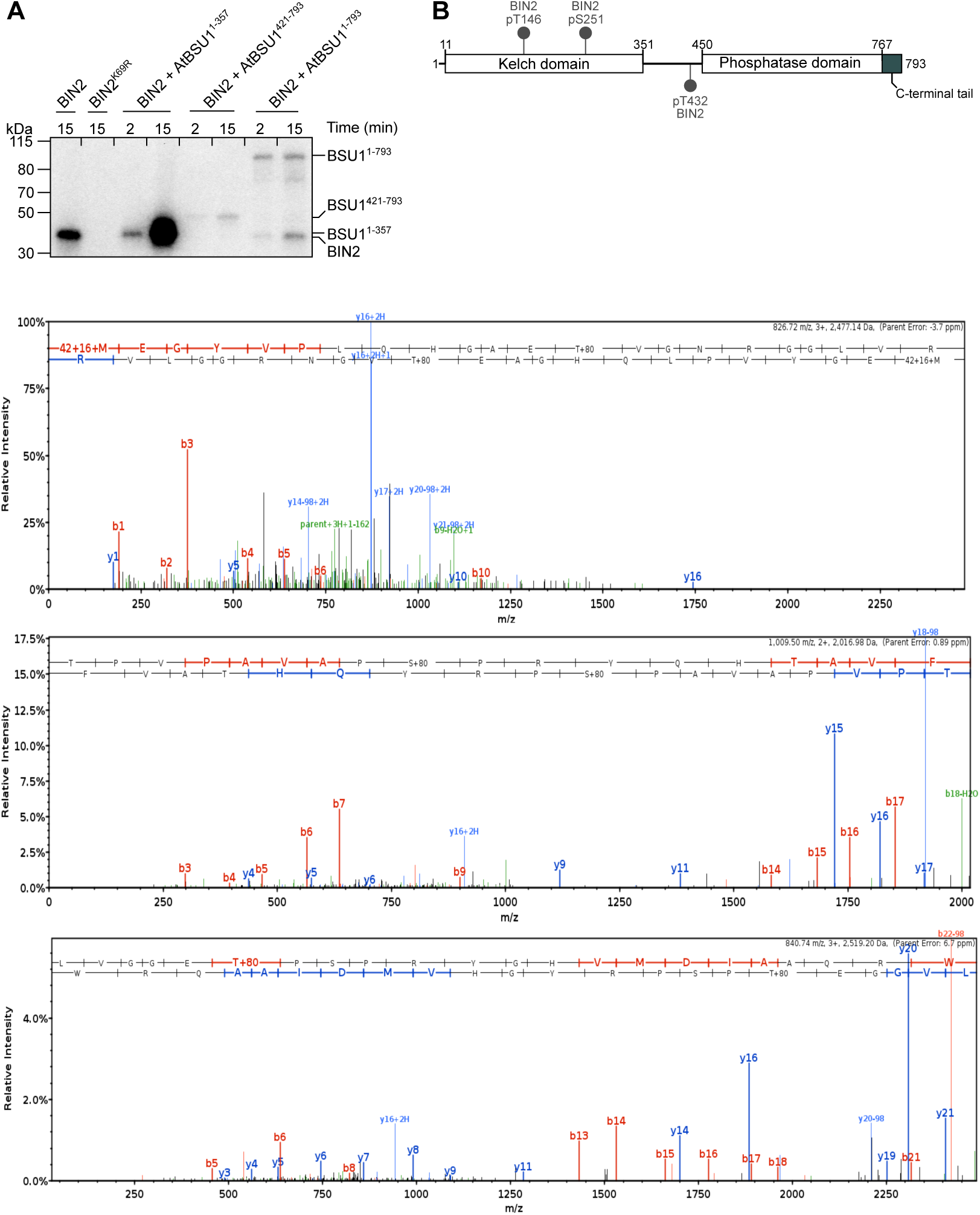
BIN2 phosphorylates BSU1 in vitro. (*A*) Autoradiograph of an in vitro kinase assay of BIN2 vs. the BSU1 Kelch domain (BSU1^1-357^), BSU1 phosphatase domain (BSU1^421-793^), or full-length BSU1 (BSU1^1-793^). BIN2^K69R^ is a kinase-dead mutant (41). (*B*) Schematic representation of BSU1, depicting the Kelch domain (residues 11–351), phosphatase domain (residues 450–767), and C-terminal tail (residues 768–793). Structured regions are shown as blocks, and putative linker regions (residues 1–10 and 352–449) as solid lines. Phosphorylated residues targeted by BIN2 are shown in gray. Below, the MS1 spectra of the phosphorylated BSU1^243-260^, BSU1^421-441^, and BSU1^141-162^ peptides are shown from top to bottom.

**Table S1.** Crystallographic data collection and refinement statistics.

|  | **BSU1<sup>1-357</sup>** | **BSU1<sup>421-793</sup>** | **BSU1<sup>421-793</sup>** | **BSU1<sup>421-793 D510N H707K</sup>** |
| --- | --- | --- | --- | --- |
|  | **Kelch** | **SAD** | **native** | **phosphatase-dead** |
| **PDB-ID** | pdb_00009TP5 |  | pdb_00009TP7 | pdb_00009TP8 |
| **Data collection** |  |  |  |  |
| Wavelength (Å) | 0.999874 | 1.282151 | 1.000029 | 1.000031 |
| Space group | *P* 4<sub>3</sub> 2<sub>1</sub> 2 | *C* 2 | *C* 2 | *C* 2 |
| Cell dimensions |  |  |  |  |
| *a*, *b*, *c* (Å) | 77.31, 77.31, 134.32 | 226.07, 97.64, 157.79 | 226.41, 97.79, 157.80 | 139.76, 44.05, 79.24 |
| $\alpha$, $\beta$, $\gamma$ (°) | 90, 90, 90 | 90, 129.23, 90 | 90, 129.41, 90 | 90, 121.07, 90 |
| Resolution (Å) | 42.40 – 2.25 (2.39 – 2.25) | 45.69 – 2.70 (2.77 – 2.70) | 45.38 – 2.10 (2.22 – 2.10) | 41.34 – 1.60 (1.69 – 1.60) |
| *R*<sub>meas</sub><sup>#</sup> | 0.225 (3.0) | 0.196 (2.93) | 0.147 (1.87) | 0.079 (1.37) |
| CC(1/2)<sup>#</sup> | 1.0 (0.30) | 1.0 (0.59) | 1.0 (0.56) | 1.0 (0.50) |
| *I*/$\sigma$ *I*<sup>#</sup> | 12.72 (1.0) | 9.4 (1.0) | 11.44 (1.20) | 11.7 (1.1) |
| Completeness (%)<sup>#</sup> | 100 (100) | 99.6 (99.6) | 99.5 (99.2) | 99.0 (98.6) |
| Redundancy<sup>#</sup> | 13.9 (13.5) | 10.4 (10.6) | 6.8 (6.8) | 3.4 (3.3) |
| Wilson B-factor<sup>#</sup> | 51.3 | 65.1 | 43.1 | 29.5 |
| **Phasing** |  |  |  |  |
| No. of Zn<sup>2+</sup> sites |  | 8 |  |  |
| Figure of merit<sup>^</sup> |  | 0.39 / 0.82 |  |  |
| **Refinement** |  |  |  |  |
| Resolution (Å) | 42.40 – 2.25 |  | 45.38 – 2.10 | 41.34 – 1.60 |
| No. reflections | 36,873 |  | 155,227 | 106,610 |
| *R*<sub>work</sub> / *R*<sub>free</sub><sup>*</sup> | 0.22 / 0.25 |  | 0.20 / 0.23 | 0.16 / 0.18 |
| No. atoms |  |  |  |  |
| protein | 4,894 |  | 32,221 | 5,029 |
| solvent | 43 |  | 870 | 139 |
| ligand | 44 |  | 68 | 77 |
| Res. B-factors<sup>*</sup> |  |  |  |  |
| protein | 70.1 |  | 58.7 | 33.43 |
| solvent | 49.2 |  | 49.0 | 38.3 |
| ligand | 76.9 |  | 76.5 | 54.6 |
| R.m.s deviations<sup>*</sup> |  |  |  |  |
| bond lengths (Å) | 0.002 |  | 0.002 | 0.015 |
| bond angles (°) | 0.49 |  | 0.53 | 1.44 |
| Ramachandran plot<sup>\$</sup>: | | | | |
| most favored regions (%) | 96.62 |  | 95.58 | 96.09 |
| outliers (%) | 0 |  | 0 | 0 |
| MolProbity score<sup>\$</sup> | 1.13 | | 1.51 | 1.64 |

<sup>#</sup>as defined in XDS (144)
<sup>^</sup>FOM before and after NCS averaging and density modification with phenix.resolve (145, 111)
<sup>\*</sup>as defined in phenix.refine (107)
<sup>\$</sup>as defined in Molprobity (114)

## Notes

### Competing Interest Statement

The authors have declared no competing interest.

### Summary of Updates

The manuscript has been revised taking into consideration the comments associated with the preprint on biorxiv. New data are shown in revised Fig. 1G, J, K and L, and SI Appendix Fig. S1. The results and disucssion sections have been revised accordingly.

